# Overlapping transcriptional programs promote survival and axonal regeneration of injured retinal ganglion cells

**DOI:** 10.1101/2022.01.19.476970

**Authors:** Anne Jacobi, Nicholas M Tran, Wenjun Yan, Inbal Benhar, Feng Tian, Rebecca Schaffer, Zhigang He, Joshua Sanes

**Author notes:** These authors contributed equally to this work (Anne Jacobi, Nicholas M Tran, Wenjun Yan).

## Abstract

Injured neurons in the adult mammalian central nervous system often die and seldom regenerate axons. To uncover transcriptional pathways that could ameliorate these disappointing responses we analyzed three interventions that increase survival and regeneration of mouse retinal ganglion cells (RGCs) following optic nerve crush (ONC) injury, albeit not to a clinically useful extent. We assessed gene expression in each of 46 RGC types by single cell transcriptomics following ONC and treatment. We also compared RGCs that regenerated to those that survived but did not regenerate. Each intervention enhanced survival of most RGC types, but type-independent axon regeneration required manipulation of multiple pathways. Distinct computational methods converged on separate sets of genes selectively expressed by RGCs likely to be dying, surviving, or regenerating. Overexpression of genes associated with the regeneration program enhanced axon regeneration *in vivo*, indicating that mechanistic analysis can be used to identify novel methods for promoting regeneration of injured neurons.

## INTRODUCTION

Damage to the axons of central nervous system (CNS) neurons usually leads to permanent functional deficits. Adult CNS neurons have limited capacity to regenerate axons and form new synapses, and in many cases, they die. As Ramón y Cajal wrote a century ago, “In the adult centers, the nerve paths are something fixed, ended, and immutable. Everything may die, nothing may be regenerated. It is for the science of the future to change, if possible, this harsh decree.” (Ramon y Cajal, 1928). In an attempt to meet this challenge, many groups have used models of axonal injury to seek molecules that improve axonal regeneration and neuronal survival. One intensively studied model is optic nerve crush (ONC), which severs the axons of retinal ganglion cells (RGCs), the projection neurons that transmit visual information from the retina to the rest of the brain. In mice, ∼90% of RGCs die during the month following ONC, and few if any of the survivors extend new axons more than a few hundred micrometers past the site of injury (Winter et al., 2021). Using this model, several interventions have been discovered that improve survival and/or axon regeneration, but to date none has been sufficient to restore useful vision.

Our goal in this study was to analyze the molecular effects of these interventions, with the aim of elucidating pathways that promote or constrain neuronal survival and axonal regeneration. To this end, we focused on three manipulations: deletion of genes encoding PTEN (Phosphatase and tensin homolog) and SOCS3 (Suppressor of cytokine signaling 3), and delivery of ciliary neurotrophic factor (CNTF). Each of the three, separately and in combination, have been shown to enhance RGC survival and axon regeneration, with the combination of the three being more effective than any one alone (Luo and Park, 2012; Park et al., 2008; Pernet et al., 2013; Smith et al., 2009; Sun et al., 2011; Williams et al., 2020; Xie et al., 2021).

Our strategy relied on high-throughput single cell RNA-sequencing (scRNA-seq). We profiled uninjured, injured, and treated RGCs and classified them into 46 distinct types using recently described criteria (Tran et al., 2019). We showed previously that survival varies nearly 100-fold among types following ONC (Duan et al., 2015; Tran et al., 2019), so we asked whether interventions that enhance survival act selectively on vulnerable or resilient types. Second, we collected RGCs that had regenerated axons and compared them to RGCs that had survived but not regenerated, asking whether interventions promote regeneration of particular types. Next, we used five independent computational methods to analyze gene expression by RGC type, state, time after injury, and intervention, seeking expression patterns that correlated with any of these variables. These methods converged on three groups of genes, one preferentially expressed by RGCs destined to die, another by RGCs that survived but did not regenerate, and a third by RGCs that regenerated axons. These expression modules provide insights into molecular mechanisms that regulate neuronal survival and axon regrowth. Finally, we showed that manipulating several genes from the regeneration module enhances axonal regeneration following ONC, supporting the idea that this strategy can serve as a novel source of therapeutic targets. In a companion paper, we provide further computational and functional analyses of genes that play key roles in the survival of injured RGCs (Tian et al., accompanying paper).

## RESULTS

We analyzed effects of three manipulations known to promote survival and axon regeneration from RGCs following ONC: deletion of Pten, deletion of Socs3, and overexpression of CNTF. PTEN and SOCS3 are endogenous inhibitors of mTOR and Jak/Stat signaling, respectively, and CNTF activates Jak/Stat signaling. As detailed in the Discussion, their modes of action in promoting axon regeneration are incompletely understood. We tested them in three combinations: (1) conditional deletion of Pten (P_CKO_), (2) conditional deletion of Pten combined with over-expression of CNTF (C/P_CKO_) and (3) conditional deletion of both Pten and Socs3 combined with over-expression of CNTF (C/PS_CKO_). Mice also bore the Thy1-YFP line 17 transgene (called YFP17 here; (Feng et al., 2000; Sun et al., 2011), which selectively labels RGCs.

Experiments were initiated by intravitreal injection of AAV2-Cre alone or in combination with AAV2-CNTF in YFP17^het^Pten^flox/flox^ or YFP17^het^Pten^flox/flox^;Socs3^flox/flox^ mice. With this route of administration, the AAV2 serotype primarily infects cells of the ganglion cell layer, which contains RGCs and displaced amacrine cells, so deletion is strongly biased to these two cell types. Two weeks following injection, we collected retinas from some of the treated mice and performed ONC on others, then collected retinas 2, 7, or 21 days later (Figure 1A). We also collected retinas from AAV2-Cre-infected YFP17^het^ mice, and uninfected YFP17^het^Pten^flox/flox^ or YFP17^het^Pten^lox/flox^;Socs3^flox/flox^ mice (Table S1). In subsequent analyses we found no significant differences in types, type frequencies or gene expression among these latter groups; we therefore pooled data from them and refer to the combined group as !wild-type” (WT) hereafter.

**Figure 1:**
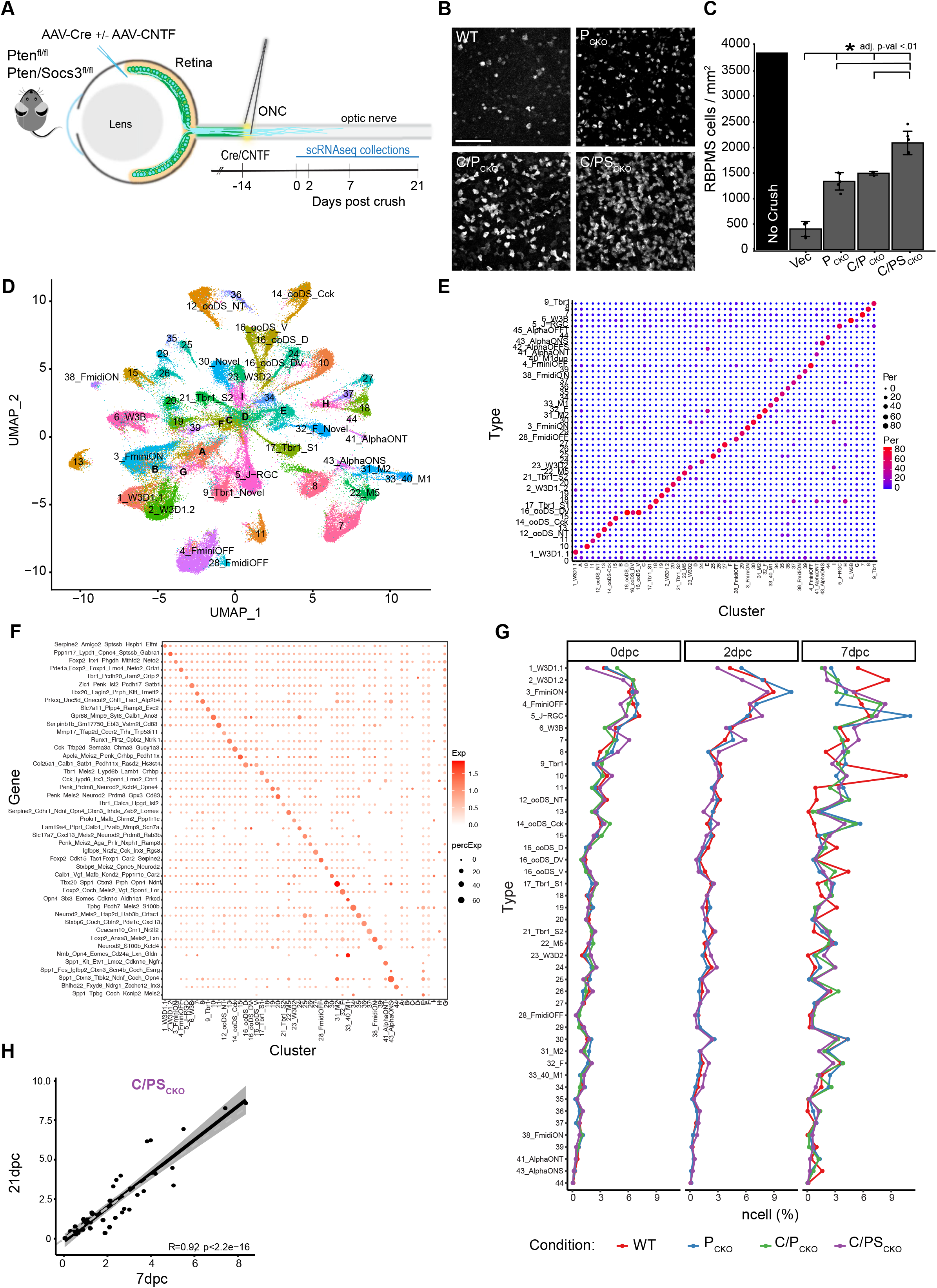
Interventions preserve type identity and increase survival of most RGC types after ONC. **A)** AAV2-Cre was injected into the vitreous body of PTEN^f/f^ (P_CKO_) or Pten^f/f^Socs3^f/f^ (PS_CKO_) mice to delete the floxed genes 2 weeks before crushing the optic nerve (ONC). AAV2 encoding ciliary neurotrophic factor (CNTF) was co-injected as indicated (C/P_CKO_ or C/PS_CKO_). RGCs were collected for scRNA-seq at indicated times thereafter. **B)** Immunohistochemistry in retinal whole-mounts for the pan-RGC marker, RBPMS, shows increased survival of RGCs at 21dpc following P_CKO_, C/P_CKO_ and C/PS_CKO_. Scale bar = 100µm **C)** RGC density (RBPMS+/mm^2^ cells, *adjusted p-value <0.01) at 21dpc compared to un-crushed control, measured from images such as those in **B**. **D)** scRNA-Seq data from all RGCs analyzed in this study displayed as a UMAP. Numbers indicate RGC ‘Novel’ types as defined in the atlas presented in Tran, et al. (2019). Letters (A-G) show clusters that could not be assigned to a type. **E)** Comparison of cell type mapping in the current dataset to the RGC atlas from Tran et al., (2019) shown as a confusion matrix. Dot sizes and colors represent the percentage of cells in each cluster on the x-axis that match the atlas types on the y-axis. **F)** Expression of gene marker combinations from the control RGC atlas in the current dataset. Color of the dot represents the average expression of the gene marker combination, and the dot size represents the proportion of cells expressing these markers. **G)** Proportion of types in WT and each intervention at 0, 2 and 7dpc. **H)** Scatterplot showing correspondence (R_Pearson_ = 0.92) between frequencies of C/PS_CKO_ RGCs of at 7 and 21dpc. Each dot shows one RGC type. The dark line shows best fit with confidence interval indicated in grey.

In each case, we collected RGCs by fluorescence-activated cell sorting (FACS) and profiled them by droplet-based scRNA-seq (10X platform; (Zheng et al., 2017). We identified RGCs based on their expression of pan-RGC markers including the RNA-binding protein, RBPMS, the class 4 POU-domain transcription factors, POU4F1-3 (Brn3a-c), and the glutamate transporter SLC17A6 (VGLUT2). RGCs comprised ∼90% of profiled cells. We considered only RGCs hereafter, excluding other cell classes and putative doublets. To estimate the fraction of RGCs that had been infected with AAV2 Cre and/or CNTF, we quantified sequencing reads that mapped to the WPRE element contained in our AAV vectors (see Methods). Such reads were detected in ∼80% in all libraries, with no substantial differences among RGC types (Figure S1A). *In situ* hybridization confirmed deletion of Pten from AAV2-Cre-infected P_CKO_ retina (Figure S1B).

### Type-independent enhancement of RGC survival

RGC survival was enhanced by all three interventions in the order C/PS_CKO_ > C/P_CKO_ > P_CKO_ > WT, with C/PS_CKO_ preserving >50% of RGCs at 21 days post crush (dpc) (Figure 1B,C). To ask whether these interventions selectively affect specific RGC types or evenly scale across all 46 types, we first clustered RGCs and assigned clusters to atlas types using markers we had identified and validated in wild-type mice (Tran et al., 2019). Most clusters had 1:1 matches with atlas types and >86% of RGCs could be confidently assigned to a type (Figure 1D-F; unassigned cells are discussed further below). Moreover, the specificity of type marker expression was largely retained after crush, though expression was somewhat degraded in WT 7dpc RGCs (Figure S1C). Thus, neither ONC nor interventions had detectable effects on cell type identity.

We then assessed the frequencies of RGC types at each time point and in each condition. Frequencies of RGC types did not differ significantly among groups at 0dpc or 2dpc but some modest differences emerged at 7dpc (Figure 1G). For example, two types of ON-OFF direction selective RGCs (ooDSGCs) and J-RGCs survived disproportionately in some or all groups (Figure S1D, S1E). We also grouped RGCs into subclasses (as defined in (Tran et al., 2019) to minimize variability owing to the small numbers of RGCs in some clusters. Again, frequencies were similar across interventions (r^2^=0.70-0.84), albeit with some exceptions, including disproportionate survival of Cartpt-RGCs (which include ooDSGCs) and T-RGCs (which include J-RGCs) and disproportionate loss of T5-RGCs (Figure S1F-H). Finally, we compared frequency distributions at 7dpc and 21dpc in the C/PS_CKO_ group, to ask whether some types were selectively preserved at later times but saw minimal differences (r=0.92; Figure 1H). Taken together, these data show that the improved survival driven by these interventions is observed across most RGC types and that the degree of neuroprotection generally scales with the cell type’s innate resilience.

### Overcoming type-dependent RGC axon regeneration

We next asked which RGC types regenerated axons following ONC. We first quantified the extent of regeneration by injecting a fluorophore-tagged anterograde tracer intravitreally at 19dpc, then fixing and clearing whole optic nerves at 21dpc. Regeneration was minimal in wild-type mice, but enhanced by all three interventions in the order C/PS_CKO_ > C/P_CKO_ > P_CKO_ (Figure 2A,B) (Duan et al., 2015; Park et al., 2008; Sun et al., 2011). This order was the same as that seen for survival (Figure 1B,C), but was not a consequence of enhanced survival in that the increase in regeneration (>100-fold) was far greater than the increase in survival (2-5-fold).

**Figure 2:**
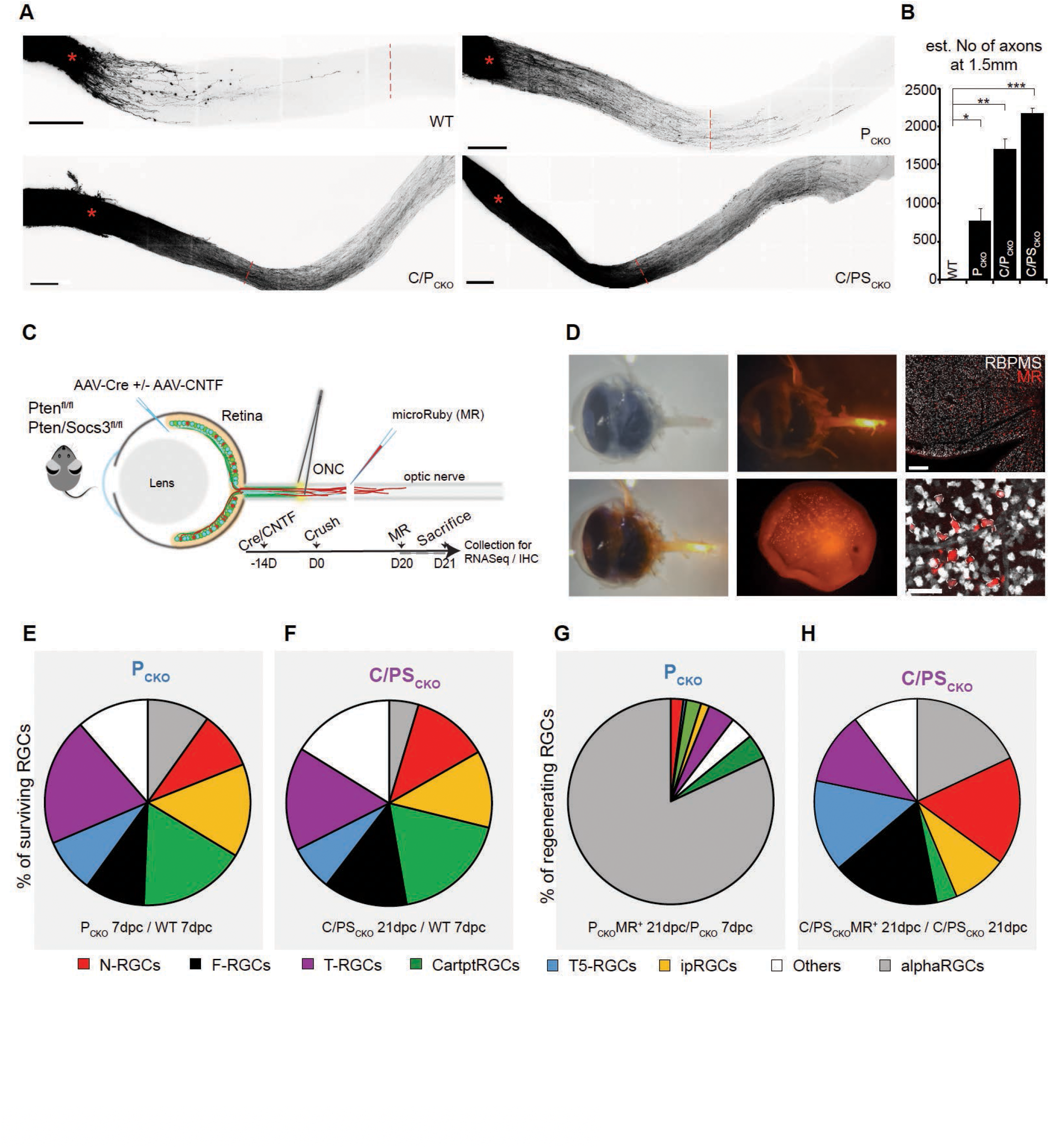
Type-independent axon regeneration in C/PS_CKO_. **A)** Maximum projections through cleared optic nerves showing anterograde-labeled RGC axons at 21dpc following injection of AAVs encoding Cre and/or CNTF into Pten^fl/fl^ or Pten^fl/fl^ Socs3^fl/fl^ mice. An empty vector was injected into WT mice. Scale bar: 250µm; X = crush site; red dashed lines are 1.5mm from X. **B)** Numbers of regenerating axons 1.5mm distal to injury site at 21dpc, from images such as those shown in **A**. Error bar: SEM; p value Students t test: * < 0.05; ** <0.01; *** < 0.001. Vector n = 6, P_CKO_ n = 5, C/P_CKO_ n = 4, C/PS_CKO_ n = 4. **C)** Protocol for retrograde labeling of regenerating RGCs for SS2 collection. 5% Dextran micro-Ruby (MR) was injected into the nerve stump at ∼1mm distal to ONC at 20dpc. **D)** Eyes collected 21dpc. Left and upper middle panels show fluorescence of injected MR in nerve stump. Lower middle panel shows retrogradely labeled RGCs in a dissected C/PS_CKO_ retina. Right panels from retina as in D but labeled with anti-RBPMS (pan-RGC marker in grey). Dashed lines outline MR^+^ RGCs. Scalebar = 500µm (top), 50µm (bottom). **E)** and **F)** Proportions of surviving RGCs by subclass in P_CKO_ (**E**) and C/PS_CKO_ (**F**) among RGCs collected by the 10x Genomics platform. All subclasses are present among surviving RGCs. **G)** and **H)** Proportions of regenerating RGCs (MR^+^) among retrograde-labeled RGCs collected by SS2 shown by subclass in P_CKO_ (**G**) and C/PS_CKO_. (**H**) Most (83%) regenerating P_CKO_ RGCs are alphaRGCs but in C/PS_CKO_ all subclasses regenerated in approximate proportion to their frequency among survivors.

To separate RGCs that regenerated from those that survived but did not regenerate, we used a retrograde labeling method in which we injected a small fluorescently labeled dextran (micro-Ruby, MR) into the nerve stump ∼1.5mm distal to the crush site at 20dpc (Zhang et al., 2019). The dextran was taken up by regenerating axons and retrogradely transported to RGC somata, which were also labeled with YFP (Figure 2C,D). Control experiments demonstrated that the method was efficient and specific: most RGCs were labeled in uninjured retina but few if any RGCs were labeled following ONC in wild-type retina (Figure S2A). Thus, this labeling strategy efficiently and specifically marked regenerating axons that, based on dye spread, had extended ≥1mm from the crush site (Figure 2D). In contrast, non-retrogradely labeled RGCs were highly enriched for those that regenerated minimally. We used FACS to isolate regenerating (MR^+^YFP^+^) and non-regenerating (MR^-^YFP^+^) RGCs 24hrs after tracer injection (Figure S2B), collected single cells in individual wells, and performed scRNA-seq using SmartSeq2 (SS2).

We obtained 120 single RGC transcriptomes from P_CKO_ retinas (all MR^+^) and 245 from C/PS_CKO_ retinas (179 MR^+^ and 66 MR^-^). The distribution of RGCs that had regenerated axons differed dramatically between the two groups: Pten deletion selectively promoted regeneration of alphaRGCs (Figure 2G), consistent with our previous results (Duan et al., 2015), whereas frequencies of regenerating RGCs in the C/PS_CKO_ group approximately mirrored their proportion among survivors (Figure 2F, H).

Combining measurements of the amount of regeneration promoted by P_CKO_ and C/PS_CKO_ (Figure 2B) and the RGC types that regenerate (Figure 2G,H) revealed that similar numbers of regenerating RGCs are alphaRGCs in P_CKO_ and C/PS_CKO_ mice, indicating that most of the “additional” regenerating RGCs in the latter are non-alphaRGCs. We conclude that the combined treatment overcomes the type-specific barriers seen when only Pten is deleted.

### Injury-independent effects of Pten, Socs3 and CNTF

We next undertook a detailed analysis of gene expression changes that result from manipulation of Pten, Socs3 and CNTF, some of which are likely to underlie their beneficial effects. Because we introduced AAV2 vectors two weeks prior to ONC to ensure efficient expression, it was possible that some transcriptional changes preceded injury. This might be akin to the !conditioning effect” observed in dorsal root ganglia, where a !priming” injury to the peripheral branch of the sensory neurons induces growth-promoting transcriptional changes that enhance regeneration of axons following a later injury to the central branch (Neumann and Woolf, 1999; Richardson and Issa, 1984). Therefore, we began by assessing injury-independent effects of these manipulations.

Two weeks after AAV2 injection (i.e., 0dpc), 74 genes were up-regulated in C/P_CKO_ and 51 in C/PS_CKO_ RGCs compared to WT (>1.5-fold, Figure 3A and Table S2). Several of these genes have been annotated as regeneration-associated genes (RAGs) in prior studies (e.g., *Bdnf, Stat3, Tubb3*; Figure 3A, S3A) (Chandran et al., 2016; Renthal et al., 2020; Yang et al., 2020). For a more comprehensive view, we generated a !RAG score” composed of the 306 differentially expressed (DE) genes selectively expressed by regenerating (retrogradely labeled) RGCs in the C/PS_CKO_ condition (see below, Figure 4). This score was substantially higher in C/P_CKO_ and C/PS_CKO_ RGCs compared to P_CKO_ and WT RGCs (Figure 3B).

**Figure 3:**
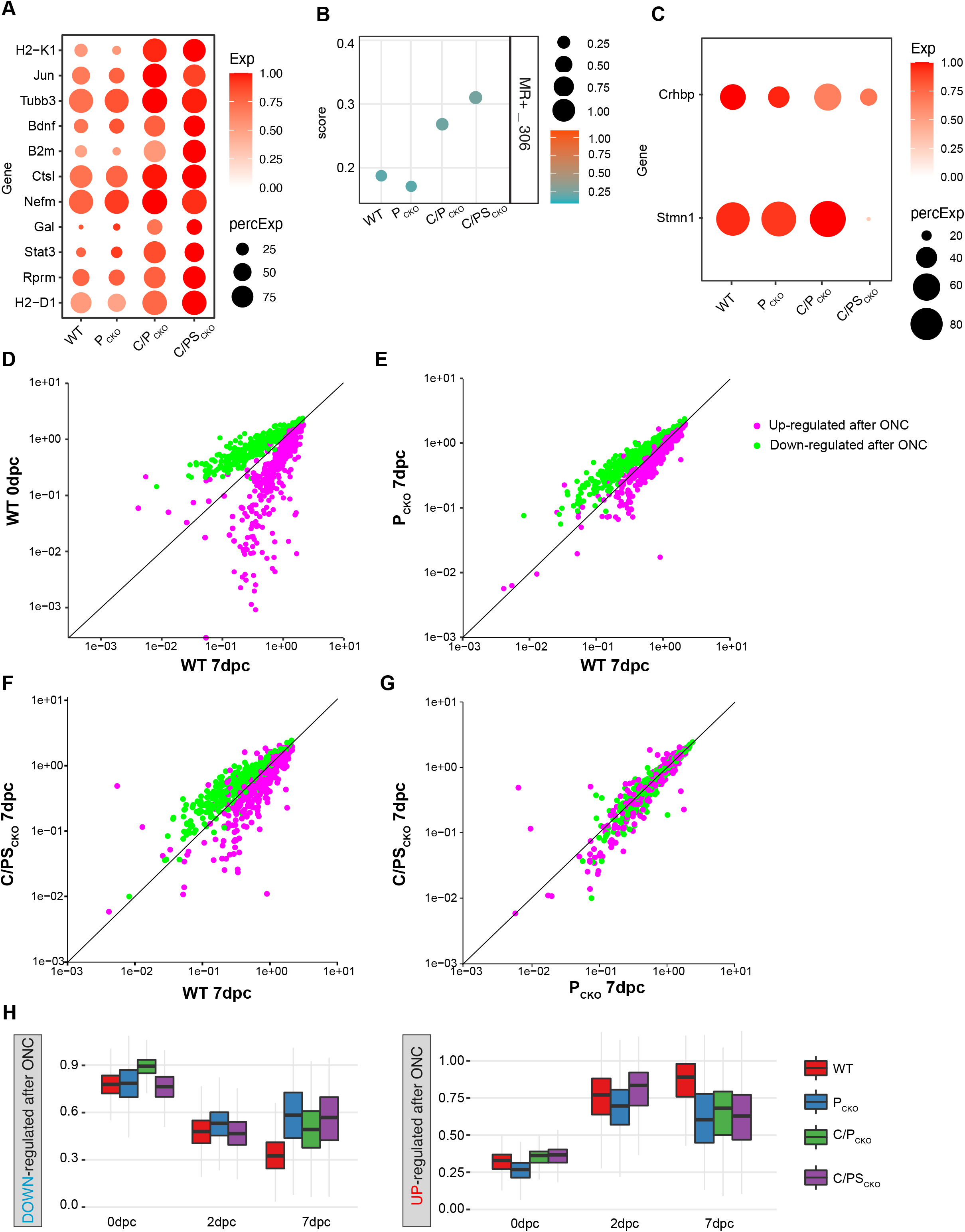
Injury independent effects and mitigation of injury response induced by interventions. **A)** Dotplot showing genes upregulated in C/P_CKO_ and C/PS_CKO_ compared to WT and PCKO prior to injury (0dpc, 2 weeks after AAV injection). **B)** !Composite RAG score” (defined in text), compiled from 306 genes selectively expressed in regenerating (MR^+^) C/PS_CKO_ RGCs. **C)** Dotplot showing Crhbp and Stmn1 downregulation in C/PS_CKO_ prior to injury. **D-G)** Scatterplots showing expression level in current dataset of 771 genes identified as upregulated (pink) or downregulated (green) after ONC (0.5-14dpc) in a previous study (Tran et al. 2019). Responses in current data (WT 7dpc vs 0dpc) are similar to those in Tran et al., showing reproducibility (**D**). In P_CKO_, (**E**) or C/PS_CKO_ (**F**), expression changes are attenuated or reversed. There are only minor differences between overall expression between P_CKO_ and C/PS_CKO_ **(G)**. **H)** Boxplots of composite scores showing average expression of the up- or downregulated genes from **(D)-(F)** at 0, 2 and 7dpc. Horizontal line = mean, box below = 25^th^ percentile, box above = 75^th^ percentile, grey lines = whiskers / range.

**Figure 4:**
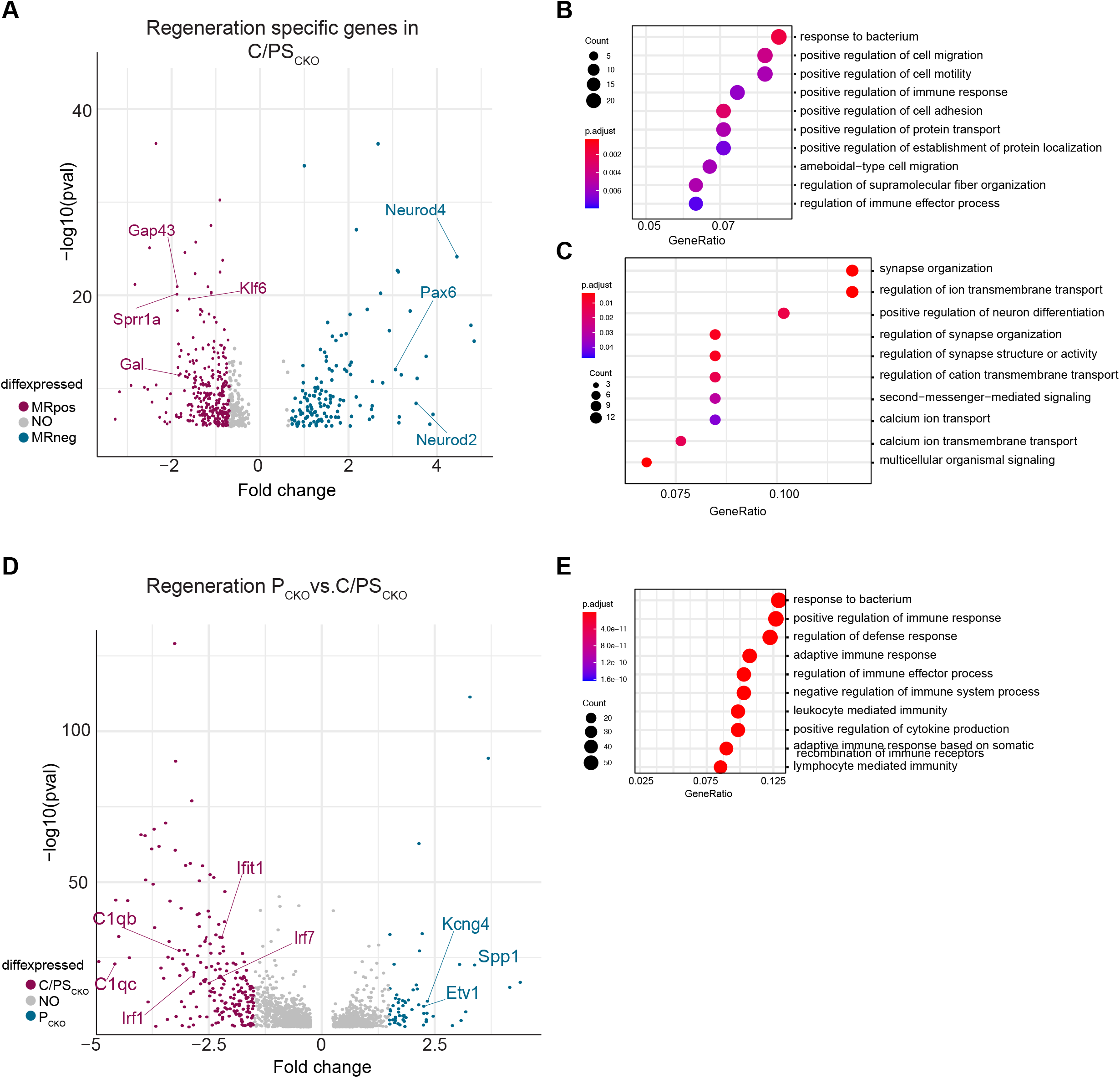
Gene expression analysis of regenerating RGCs. **A)** Volcano Plot of genes differentially expressed between MR^+^ and MR^-^ RGCs from C/PS_CKO_ retinas at 21dpc. p-value < 0.05, logFC> 0.7. Grey dots are genes not considered significantly DE. **B, C)** Dotplots highlighting Top10 GO-pathways enriched in regenerating (MR^+^) RGCs compared to surviving (MR^-^) RGCs (**B**) or in surviving compared to regenerating RGCs (**C)**. **D)** Volcano Plot of genes differentially expressed between MR^+^ P_CKO_ and C/PS_CKO_ RGCs. Genes associated with immune response or alphaRGCs are indicated in red and blue, respectively. Grey dots as in **(A)**. **E)** Dotplot highlighting Top10 GO-pathways enriched in regenerating MR^+^ C/PS_CKO_ RGCs compared to MR^+^ P_CKO_ RGCs.

A smaller set of genes was downregulated in intact C/PS_CKO_ RGCs relative to controls (16 genes, >1.5-fold, FDR < 0.01). They included *Crhbp*, which inhibits RGC regeneration (Tran et al., 2019) and *Stmn1*, which destabilizes microtubules and therefore could impair axon regeneration (Rubin and Atweh, 2004) (Figure 3C, S3B). Together, these results imply that the interventions we tested may act in part by inducing an axon regeneration program prior to the injury.

### Interventions attenuate transcriptional responses of RGCs to injury

To assess gene expression changes following intervention and ONC, we first plotted genes that had been identified as being down-regulated (n= 412 - green dots) or up-regulated (n= 359 - magenta dots) in WT mice in our previous study (Tran et al., 2019). Nearly all showed the same responses in the new dataset (Figure 3D), demonstrating (a) that our transcriptomic methods were reproducible and (b) that the 2 and 7dpc time points captured both early and late-stage injury response genes from the previous time course (0.5-14dpc).

The global effect of deleting Pten was to attenuate these changes. At 7dpc, 73% of the genes downregulated after ONC in WT mice were expressed at significantly higher levels in P_CKO_ than in WT mice, and 51% of the genes upregulated after ONC in WT mice were expressed at significantly lower levels in P_CKO_ than in WT mice (Figure 3E). No down-regulated and only 6% of the upregulated genes displayed the opposite trend. Thus, Pten deletion counteracts injury-induced changes in gene expression. Surprisingly, these injury-induced changes in gene expression differed little between RGCs from P_CKO_ retina and C/PS_CKO_ retina (Figure 3E-G), suggesting that Pten is the main driver of this mitigation effect.

To determine when Pten acts, we also compared the expression of these DE genes at 0 and 2dpc. Interventions had minimal effect on changes in gene expression at either of these times but counteracted further alterations in expression patterns between 2dpc and 7dpc (Figure 3H). Thus, signaling pathways regulated by Pten deletion divert RGCs from a degenerative path a few days after injury by re-establishing a more !normal” expression program that supports survival.

### Genes selectively expressed by regenerating RGCs

Even under conditions that ensure long-term survival, most RGCs fail to regenerate axons (<10% in C/PS_CKO_ retina, calculated from Figures 1C and 2B). To identify genes that might promote regeneration, we compared regenerating (MR^+^) and non-regenerating (MR^-^) RGCs in the C/PS_CKO_ intervention as described above (Figure 2).

Regenerating RGCs selectively expressed 306 genes (>1.5 fold, FDR < 0.01), many of which have been classified as RAG genes (e.g., *Sprr1a, Klf6, Gap43*; (Bonilla et al., 2002; Jankowski et al., 2009; Latremoliere et al., 2018; Richner et al., 2014); Figure 4A). Gene ontology analysis showed enrichment of pathways related to regulation of cell migration/motility and cell adhesion (Figure 4B). In contrast, RGCs that survived but did not regenerate showed enrichment of pathways related to synapse organization and neuronal differentiation, including transcription factors implicated in neurogenesis such as *Neurod2, Neurod4* and *Pax6* (Figure 4A, C) (Table S3) (Cherry et al., 2011; Marquardt et al., 2001).

As noted above, C/PS_CKO_ leads to more robust and less type-dependent regeneration of RGCs than P_CKO_ alone. To gain insight into factors that underlie this added benefit, we compared the transcriptomes of regenerating (MR^+^) RGCs in C/PS_CKO_ and P_CKO_ retinas. Unsurprisingly, genes differentially expressed by regenerating P_CKO_ RGCs included marker genes for alphaRGCs (e.g., *Spp1, Kcng4*; Duan et al., 2015), consistent with the type-specific regeneration of this group. In contrast, pathways selectively upregulated by the triple intervention were related to immune responses, particularly interferon and cytokine signaling, rather than to specific RGC types, (Figure 4D,E). Thus, while survival appears to be promoted via the re-activation of developmental processes, axon regeneration may require the additional upregulation of RAGs and immune response programs and may contribute to overcoming the type-specific barriers of axon regeneration.

### Gene expression programs associated with degenerating, surviving, and regenerating RGCs

In each of the four conditions we analyzed (WT, P_CKO_, C/P_CKO_, and C/PS_CKO_), different proportions of RGCs degenerated, survived and regenerated axons. To identify gene expression programs involved in these three distinct responses, we performed four sets of analyses: (1) We combined all transcriptomes from each intervention at 7dpc and searched for intervention-specific pathways. (2) We used Monocle3 (Qiu et al., 2017) to analyze gene co-expression in RGCs on a single-cell level. (3) We used Seurat (Hao et al., 2021) to assess RGCs that failed to map definitively to a specific type. (4) We used Single-Cell Regulatory Network Inference and Clustering (SCENIC; Aibar et al., 2017; Van de Sande et al., 2020) to identify gene-regulatory networks (GRNs). Remarkably, all four methods converged on a common set of gene modules differentially expressed by degenerating, surviving and regenerating RGCs.

#### Intervention-dependent gene expression

We combined all RGCs subjected to each intervention and analyzed them in two ways: comparing each condition (WT, P_CKO_, C/P_CKO_, and C/PS_CKO_) at 7dpc) to the sum of the others, and each to the next less complex (P_CKO_ to WT, C/P_CKO_ to P_CKO_ and C/PS_CKO_ to C/P_CKO_). We sorted DE genes for both comparisons into 4 modules by k-means clustering (nclust = 4) (Figure 5A, B, Table S4). In principle, the first comparison would generate intervention-selective genes, while the second would reveal incremental effects of each added manipulation. In fact, however, both comparisons generated similar modules as judged by a hypergeometric test (Figure S4A).

**Figure 5:**
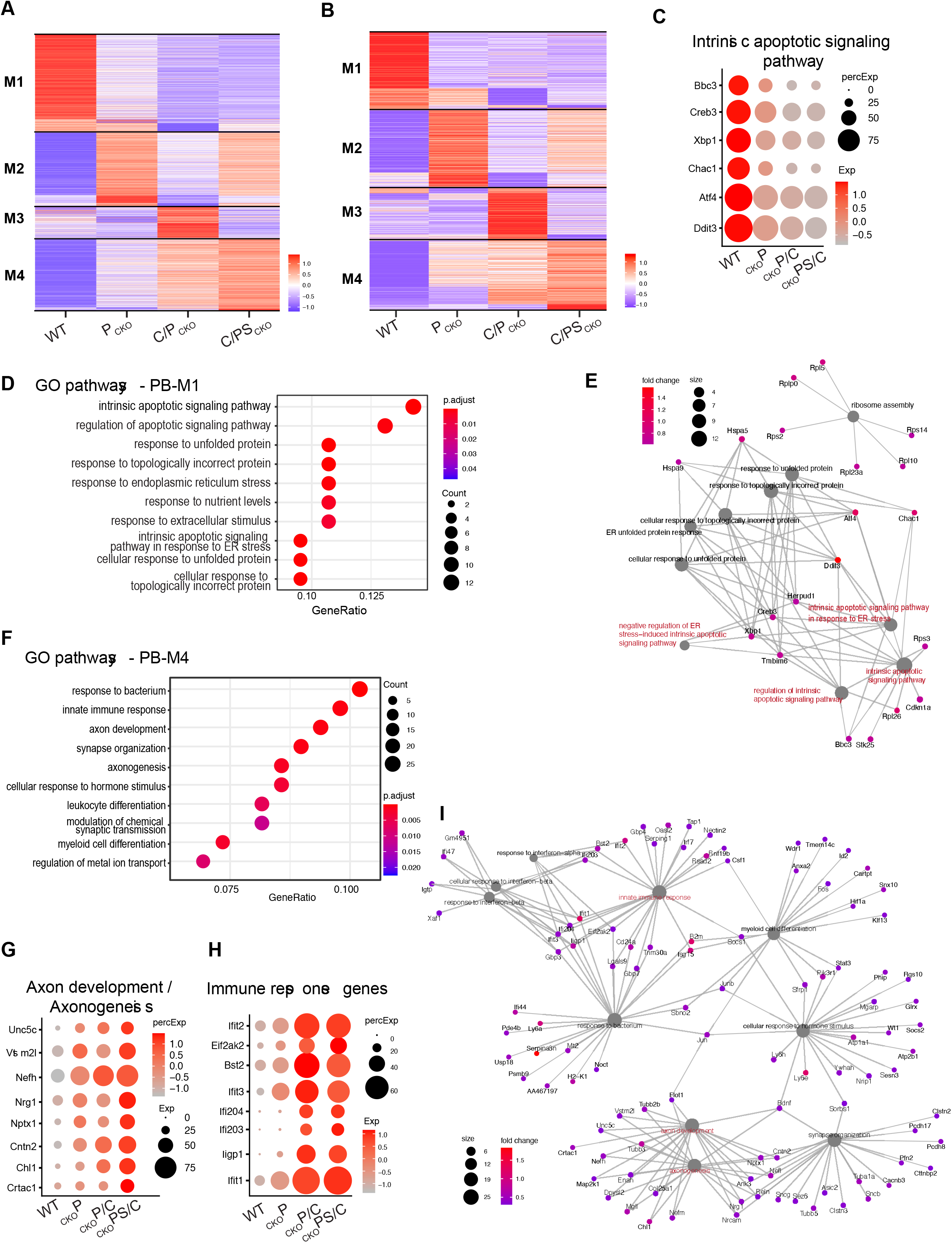
Gene modules revealed as genes selectively regulated by individual interventions. **A,B)** Heatmaps showing genes selectively expressed at 7dpc following each intervention, as calculated for each intervention against all others (**A**) or for each intervention compared to the one to its left (**B)** Expression values of each gene (row) are averaged across all RGCs in an intervention (columns) and then z-scored prior to plotting. Black bars separate genes into 4 modules (PB-M1-4). **C)** Dotplot showing expression of selected apoptotic pathway associated genes from PB-M1 at 7dpc. **D,F)** Top10 GO-pathways enriched in PB-M1 (**D**) or PB-M4 (**F**) (logFC > 0.6, FDR < 0.001) **E,I)** Cnet plot of Top10 pathways for PB-M1 **(E)** or PB-M4 **(I)** with associated genes. Color of dots represents the fold change of genes. Size of the grey dots refer to the number of genes enriched with the GO-term. **G,H)** Dotplots showing expression of genes implicated in axonogenesis (**G)** and immune responses (**H**) from PB-M4.

As assessed by gene ontology pathway (GO) analysis, genes in !pseudo-bulk” Module 1 (PB-M1) were associated with pathways related to apoptotic signaling and stress (Figure 5C-E and S4B). Many exhibited a gradual decline across interventions (WT > P_CKO_ > C/P_CKO_ > C/PS_CKO_; Figure 3C). Conversely, PB-M4 was lowest in WT and increased across treatments (WT < P_CKO_ < C/P_CKO_ < C/PS_CKO_; Figure 5G, H). Many genes in this module were associated with axon growth and development, innate immune responses, and hormone signaling (Figure 5F-I). RAGs were also prominent in this module (Fig. S4J). PB-M2 and 3 were expressed at highest levels in P_CKO_ and C/P_CKO_ respectively; enriched pathways included those related to synaptic transmission (PB-M2) and cytoplasmic translation (Figure S4C-H). Together, this comparison shows that deleting Pten from RGCs, augmenting Pten deletion with CNTF and then additionally deleting Socs3 enhances expression of genes associated with survival and regeneration (Fig. S4I) and attenuates expression of genes associated with death and degeneration.

#### Population-specific gene expression modules

To analyze gene co-expression in single cells, we reclustered RGCs at 7dpc using Monocle3 (Qiu et al., 2017). The 40 resulting clusters (Figure 6A, Figure S5A) grouped into 6 modules, MO-M1-6. Each included cells from all four conditions (Figure 6B) but occupied distinct (albeit overlapping) regions in UMAP space (Figures 6D-I). MO- M1-3 were closely related to each other (see dendrogram at left of Figure 6A) as were MO-M5 and 6.

**Figure 6:**
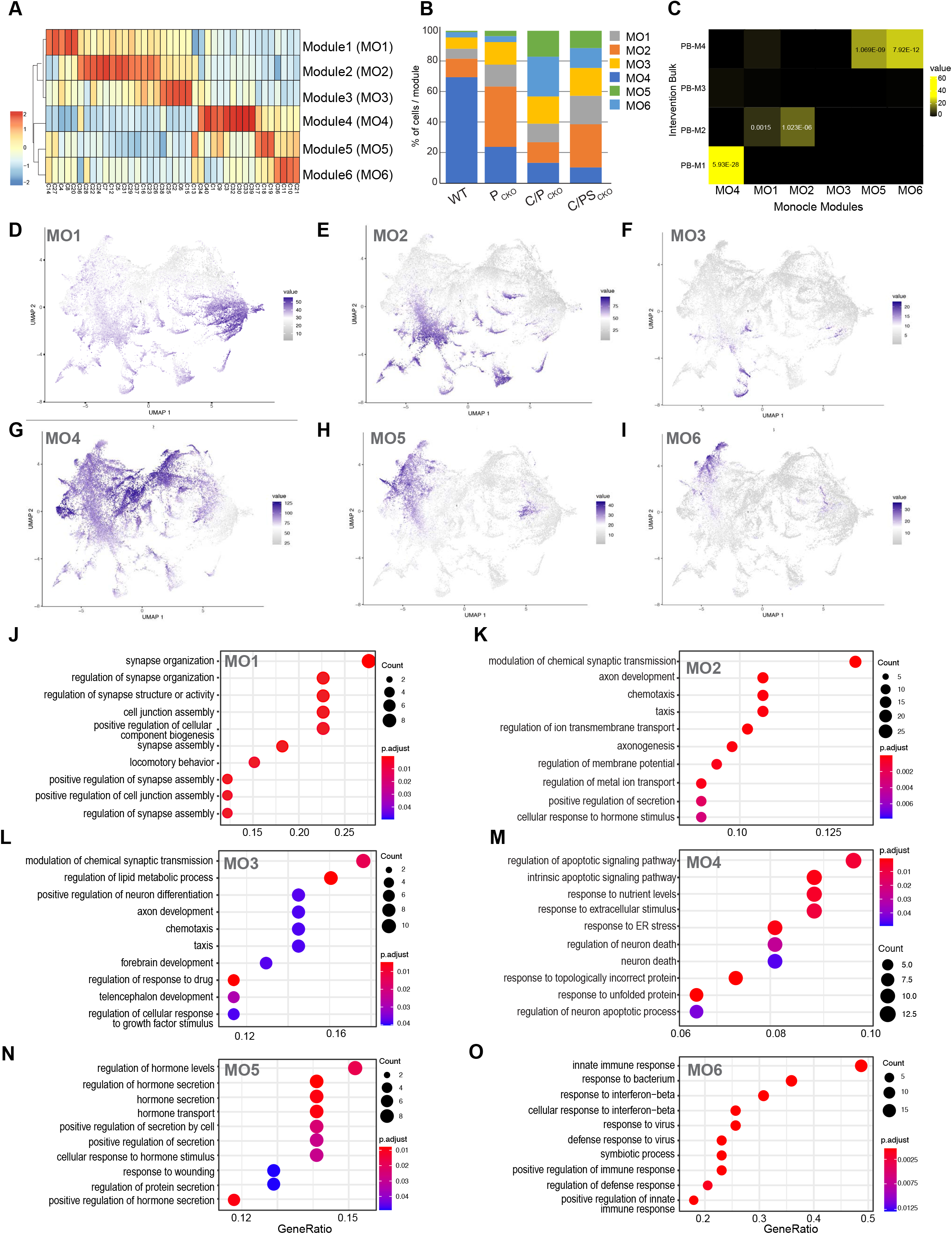
Gene modules revealed by single cell analysis using Monocle. **A)** Expression of 6 co-expression gene modules across the 40 Monocle clusters at 7dpc. Their relationships are indicated by the dendrogram to the left. Module expression is averaged across rows. **B)** Proportions of RGCs belonging to sets of Monocle clusters predominantly enriched for the indicated co-expression Module in each intervention. **C)** Heatmap of statistical enrichment using the hypergeometric test indicating correspondence between modules obtained from Pseudo-bulk analysis (Figure 5A) and Monocle analyses modules. Statistically significant p-values are shown. **D-I)** UMAPs show enrichment co-expression Modules across RGC populations. **J-O)** Top10 GO-pathways for each Monocle module. All genes contributing to the modules were considered. (logFC > 0.6, FDR < 0.001).

MO-M1-3 were enriched in pathways related to synaptic transmission (MO-M1), axonogenesis and axon development (MO-M2, Figures 6J-K S5D-F). Key genes included *Unc5d, Robo2* and *Nrgn*. MO-M3 was expressed by a subset of cells in MO-M1 and 2. These modules showed closest relation to PB-M2 from the pseudo-bulk analysis (Figure 6C). MO-M4 was strongly enriched for genes associated with intrinsic apoptotic and stress pathways (Figure 6M, S5G, Table S5), and was related to PB-M1 in the pseudo-bulk analysis (Figure S6C). MO-M5 and 6 partially overlapped in UMAP space (Figure 6H, I) and were related to PB-M4 in the pseudo-bulk analysis. Interestingly, two sets of genes present in PB-M4 were segregated into distinct modules by Monocle: genes associated with hormone and neuropeptide signaling (e.g., *Gal* and *Crh*) and axon regeneration (e.g., *Sprr1a* or *Nefh*) in MO-M5 (Figure 6N, S5H and Table S5) and genes associated with immune response and cytokine signaling (e.g., *Ifit1* and *Ifit2*) in MO-M6 (Figure 6O, S5I and Table S5). Combined immunohistochemistry and in situ hybridization confirmed that *Gal* (MO-M5) and CART (MO-M2) labeled distinct RGC populations (Figure S5B,C).

#### Injured RGCs lacking clear type identity

Although 86% of RGCs could be assigned to specific types or subclasses regardless of intervention or time after injury (Figure 1D), the remainder, comprising 7 clusters (A-G in Figure 1D) could not. To characterize these groups, we compared each of them to all other clusters. DE genes in clusters A-E were closely related to those derived from Monocle analysis: A and E resembled MO-M5 and 6, enriched in genes characteristic of regenerating cells; B resembled MO-M4, characteristic of degenerating cells; and C and D resembled MO-M1, rich in characteristics of synapse organization and transmission (Figure S6A-F). Expression patterns in the other two clusters (F,G), comprising ∼17% of this cohort (A-G) were more difficult to interpret. Thus, the unmapped clusters were largely composed of cells in which “state”- rather than type-driven expression dominated.

#### Gene regulatory networks

To seek transcription regulators of degeneration, survival and regeneration, we used SCENIC (Van de Sande et al., 2020) to identify cell-specific gene regulatory networks (GRNs), i.e., groups of transcription factors [TFs] and their predicted target genes, together called regulons (see Methods). We plotted a heatmap of the expression from the top 20 regulons at 7dpc at single cell resolution, revealing sets of cells grouped by regulon activity (Figure 7A), five of which (SC-M1-5) we highlight below.

**Figure 7:**
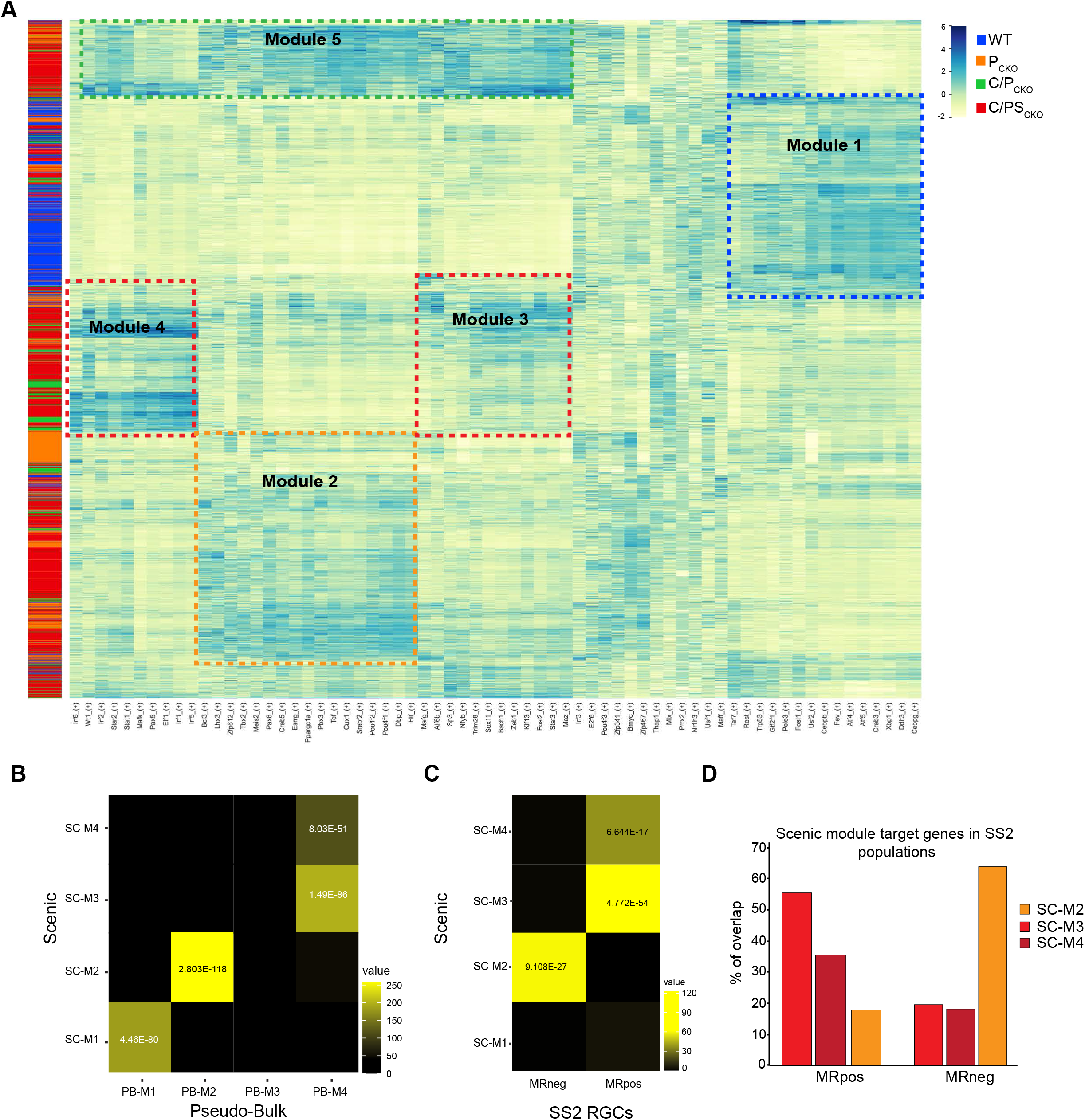
Gene regulatory modules revealed by Scenic. **A)** Heatmap of top regulon expression level (transcription factors (TF) and their putative downstream targets) in each cell at 7dpc established by Scenic analysis. Each row is a single RGC, with color bar at left indicating intervention type. Each column is a single regulon, with the TF listed at bottom. Dotted lines indicate 4 modules discussed in the text. **B,C)** Heatmaps of statistical enrichment using the hypergeometric test indicating the possibility of overlap between Scenic module regulons (TFs and potential regulated target genes; y-axis) and PB-M1-4 from Pseudo-Bulk analysis (**C**) or genes selectively expressed in regenerating (MR^+^) or surviving (MR^-^) RGCs from microRuby analysis (**D**). Statistically significant p-values are shown. **D)** Proportions of SCENIC module target genes shared with genes enriched in regenerating (MR^+^) or surviving (MR^-^) RGCs obtained from microRuby dataset.

SC-M1 was enriched for degeneration and cell death-associated TFs including *Atf4, Ddit3*, and *Cebpg*, which emerged as key promoters of degeneration in the accompanying study by Tian et al. (accompanying paper). It was related to the death-associated modules derived from pseudo- bulk (PB-M1) and Monocle (MO-M4) analyses, and a majority of its cells were from WT retinas (Figure 7A,B). SC-M2 was enriched for TFs associated with RGC differentiation and neuron development including *Pax6, Meis2*, and *Pou4f1* and *2*. It was related to the survival-associated modules PB-M2 and MO-M1-3 (Figure 7B). SC-M3 and 4 were related to the regeneration-associated modules PB-M4, MO-M5 and MO-M6. Regeneration associated genes were differentially distributed between these modules: SC-M3 was enriched for canonical RAG TFs (e.g., *Sox11, Maz* and *Stat3*), while SC-M4 included TFs associated with innate immunity and interferon signaling (e.g., *Irf1, 2,* and *8)* which were regulated in the same RGCs as MO-M6. Target genes regulated by these TFs include additional TFs implicated in immune responses (e.g., *Stat1*), axon growth (e.g., *Creb1*) and axon regeneration (e.g., *Klf6*) (Kole et al., 2020; Wang et al., 2018)(Ma et al., 2014; Romaniello et al., 2012; Sun et al., 2015) emphasizing the complex interactions among TFs in these modules (Table S6). Finally, cells in SC-M5 expressed a combination of genes from SC-M2-4. We speculate that these cells may represent RGCs with modest regenerative ability – for example, RGCs with short regenerating axons (<1 mm), which would not have been labeled by the retrograde labeling technique we used. Consistent with these assignments, target genes in SC-M2 were similar to those selectively expressed in surviving but not regenerating cells in the SS2 dataset (MR^-^) while targets genes in SC-M3 and 4 were like those selectively expressed in the regenerating (MR^+^) group (Figure 7C, D).

While each module contained cells from each intervention, we observed intervention-specific enrichment (Figure S6G), which we quantified by calculating a Regulon Specificity Score (see Methods) for all regulons in each intervention. Consistent with other analyses presented above, the highest scores in WT RGCs were associated with degeneration (with *Ddit3* as the top gene), while C/PS_CKO_ RGCs were enriched for regulons associated with regeneration (with *Wt1* and *Stat3* being the top 2 genes).

We also used SCENIC to measure gene regulatory networks at 0dpc and 2dpc. At 0dpc, the most distinctive group was related to SC-M3 and 4, consisting of RAG and interferon-beta signaling TF’s including *Stat1,2,3,5b* and *Irf1,5,7*. It was highly enriched for RGCs from the C/PS_CKO_ condition (Figure S6H), consistent with the finding that these manipulations induce a pro-regenerative “conditioning-like” effect prior to injury (Figure 3). At 2dpc, modules resembling SC-M1 (enriched for cell death related TF’s) and SC-M2 (neurodevelopment related TF’s) appeared (Figure S6I). Lastly, a group of cells did not clearly associate with a 7dpc module, instead they were enriched for TFs from multiple different states (SC-M6 in Fig. S6I), which suggests that these RGCs cells could be in a transitional state.

Taken together, our transcriptional analyses identified expression modules associated with cell states underlying degeneration, survival/initiation of regeneration, and long-distance axon regeneration. Expression modules identified by independent analyses yielded largely consistent groupings. (Degeneration: PB-M2, MO-M4, SC-M1, Seurat B; Survival: PB-M2, Monocle MO-M1/2/3, SC-M2, Seurat C, D, MR^-^; Regeneration: PB-M4, Monocle MO-M5/6, Scenic SC-M3/4, Seurat A, E, MR^+^). These results led us to test the idea that overexpression of genes prominent in the regeneration-associated modules could in fact enhance the regenerative ability of injured RGCs.

### Regeneration-associated genes promote axon regeneration

Manipulation of Pten, Socs3 and CNTF enhance RGC survival and regeneration, but have drawbacks as therapeutic targets: PTEN and SOCS3 are tumor suppressors, and recombinant CNTF shows minimal effect on its own. A main motivation of our work was the idea that genes downstream of these interventions might provide starting points for new therapeutic developments.

To test this idea, we chose 3 genes selectively expressed by RGCs in a regenerative state. Two, galanin (*Gal*), and corticotropin-releasing-hormone (*Crh*), are members of the hormone/neuropeptide signaling group that emerged from pseudo-bulk and Monocle analyses. Galanin positively affects neuronal survival and regeneration upon PNS injury (Holmes et al., 2000). CRH is a member of the corticotropin-releasing factor family, which includes another known enhancer of RGC survival and regeneration, Urocortin (Tran et al., 2019). The third candidate, *Wt1*, a TF that can act as both tumor suppressor and oncogene (Huff, 2011; Rauscher, 1993; Yang et al., 2007) directs a regulon that is specifically enriched in C/PS_CKO_ RGCs; it has not, to our knowledge, been studied in the context of axon regeneration. All three of these genes are enriched in regenerating (MR^+^) RGCs (Figure 8B). In situ hybridization confirmed expression of *Crh*, *Gal* and *Wt1* (all MO-M5) in regenerating RGCs, while *Cartpt* (MO-M2), a marker of cells that survived but did not regenerate, lacked co-labeling with regenerating RGCs (Figure S7I).

**Figure 8:**
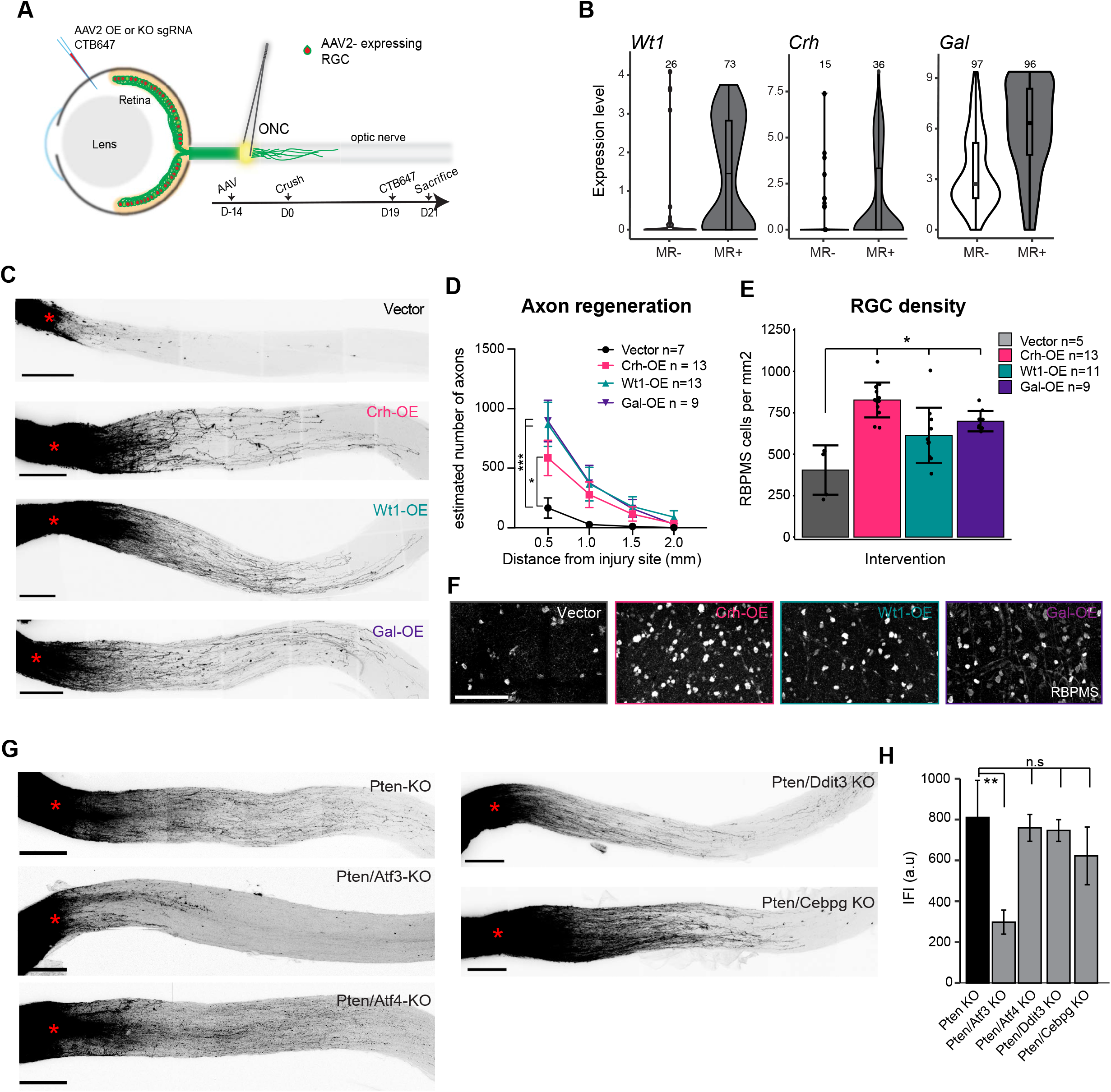
Genes affecting RCG axon regeneration. **A)** Experimental outline for *in vivo* tests of candidate regeneration-promoting genes. An AAV2 carrying a cDNA (for overexpression, OE) or sgRNA (for knockout, KO) was injected intravitreally 14 days before the crush. At 19dpc for OE or 12dpc for KO, regenerating axons were anterogradely labeled by CTB647 injection. **B)** Violin plots showing expression of OE candidates in regenerating (MR^+^) and surviving (MR^-^) RGCs. **C)** Maximum projections of cleared optic nerves showing anterograde-labeled RGC axons at 21dpc following indicated treatment. Scale bar = 250µm, red cross indicates crush site. **D)** Quantification of axon regeneration based on images such as those in **C**. Data are shown as mean %SEM. adjusted p-value * < 0.05, *** < 0.001. 2-way ANOVA or mixed effects analysis with Dunnett’s correction. **E)** RGC density (RBPMS+ cells/mm^2^; mean %SD) based on images such as those in **(F)**. *adjusted p-value <.05 (FDR). **F)** Immunohistochemistry in retinal whole mounts stained for RBPMS at 21dpc following OE-Gal, OE-Wt1 and OE-Crh. Scale bar = 100µm. **G)** Maximum projections of cleared P_CKO_ optic nerves following indicated treatment. As in **(C)** but 14dpc. **H)** Quantification of axon regeneration based on images such as those in (**G)**. Data are shown as mean %SEM with n = 4-6 each. **p<0.01.

We also tested four additional TFs – *Atf3, Atf4, Ddit3* and *Cebpg* – based on two criteria: (1) Tian et al. (accompanying paper) showed that they inhibited neuronal survival – that is, deleting them enhanced survival – and (2) *Atf4, Ddit3* and *Cebpg* direct regulons specifically enriched in the degeneration module, which was primarily comprised of WT RGCs. Unlike the other three, *Atf3* showed expression in both degenerative and regenerative RGCs (MO-M5 and -M6, Figure S7B), suggesting a dual effect.

We used AAV2 vectors to overexpress (OE) *Gal, Crh* and *Wt1*, or to mutate (knock-out, KO) *Atf3*, *Atf4, Ddit3* and *Cebpg* by introduction of gRNAs in Cas9-expressing RGCs. We injected AAV2 intravitreally 2 weeks prior to ONC and quantified RGC axon regeneration via anterograde tracing injected 2 days prior to collection (Figure 8A). Efficiency of OE or KO was demonstrated previously (Tran et al., 2019) and repeated here for selected genes using FISH (Figure S7A). The three overexpression interventions each showed a positive effect on the number of regenerating axons out to 1.5mm from the crush site (Figure 8C, D) and also enhanced RGC survival (Figure 8E, F). Interestingly, among predicted targets of *Wt1*, nearly 20% (9/52) were membrane-associated genes implicated in axon outgrowth, such as *Chl1, Cntn2* (Katic et al., 2014; Suter et al., 2020) (Table S6), underlining its potential role as a regulator for axon regeneration. In contrast, although *Atf3, Atf4, Ddit3* and *Cebpg* all enhance survival following ONC (Tian et al.; accompanying paper), they had no effect on regeneration driven by *Pten* deletion (Figure. S7C, D), emphasizing the distinct control of these two processes.

We also tested targets in combination with P_CKO._ Neither overexpression of *Gal, Wt1* or *Crh* nor deletion of *Atf4, Ddit3* or *Cebpg* led to additional regeneration (Figure S7E-H and 8G,H). In contrast, *Atf3* knockout dramatically attenuated P_CKO_ induced axon regeneration (Figure 8H).

## DISCUSSION

Following ONC in mouse, most RGCs die and hardly any of the survivors extend their axons beyond the injury site. This model has been used to seek interventions that can enhance survival and promote regeneration, but none to date has been shown to restore useful vision (He and Jin, 2016; Williams et al., 2020). The goal of this study was to investigate ways in which three of these interventions, described below, act and to identify the core molecular programs associated with axon regeneration, with the aim of finding novel therapeutic approaches. To this end we pretreated retinas in three ways prior to ONC, then used high throughput scRNA-seq to assess their effects. Our results fall into three groups. First, we analyzed the cell type-specificity of these interventions. All RGCs are similar in many respects but can be divided into ∼46 types based on morphological, physiological and molecular differences. We and others have shown that the extent of survival without intervention varies dramatically among these 46 RGC types (Bray et al., 2019; Duan et al., 2015; Tran et al., 2019). Here, we asked whether the interventions selectively affect some of them or whether their benefits are equally distributed across types. Second, we analyzed gene expression programs activated or repressed by the interventions. Finally, we tested genes identified through expression analysis by overexpression or deletion in vivo and found that some indeed promoted regeneration.

### Role of Pten, Socs3 and CNTF in neuroprotection and axon regeneration

The interventions we used were conditional deletion of the cytoplasmic phosphatase Pten, conditional deletion of the negative regulator cytokine signaling Socs3, and overexpression of the cytokine and neurotrophic factor CNTF. All interventions used AAV2 vectors injected intravitreally to manipulate target gene expression in a relatively RGC-selective manner.

PTEN antagonizes PI3-kinase by dephosphorylating lipid substrates of PI3-kinase. A major consequence of Pten deletion is the activation of a protein kinase, AKT, which in turn activates mTOR (Jaworski and Sheng, 2006; Nieuwenhuis and Eva, 2022). Earlier studies showing that Pten deletion enhanced RGC survival and regeneration provided evidence that mTOR activation is required for the effect (Park et al., 2008; Sun et al., 2011), but they did not address whether it is sufficient. PTEN and AKT also modulate other pathways (Hill and Wu, 2009; Manning and Cantley, 2007; Morgan-Warren et al., 2013) and may act in the nucleus as well as cytoplasmically (Planchon et al., 2008). The role of PTEN in axon regeneration likely involves some of these additional pathways, as interventions that more selectively activate mTOR are less effective in eliciting regeneration than Pten deletion (Park et al., 2008; Duan et al., 2015).

The second intervention was to deliver ciliary neurotrophic factor (CNTF) in parallel with Pten deletion. Addition of CNTF significantly increased RGC axon regeneration (Figure 2A, B). CNTF is a cytokine that acts in part through JAK/STAT signaling (Peterson et al., 2000) and has been shown to be a potent neurotrophic factor in multiple contexts (Fudalej et al., 2021; Richardson, 1994). It can also act in part cell-non-autonomously in retinal glial and immune cells (Müller et al., 2007; Xie et al., 2021). We induced CNTF-expression selectively in neurons of the ganglion cell layer (RGCs and amacrine cells) using the AAV2/2 serotype, and the CNTF receptor (*Cntfr*) is expressed by RGCs but we cannot exclude the possibility that the effects we saw involve additional cell types.

The third intervention was deletion of Socs3, in combination with Pten deletion and CNTF overexpression. This treatment further enhanced RGC axon regeneration and survival, consistent with previous studies (Sun et al 2011). SOCS3 acts in part as an inhibitor of the JAK/STAT signaling pathway by blocking JAK2 activity (Babon et al., 2012; Kershaw et al., 2013), which in turn can decrease the responsiveness of neurons to injury-induced cytokine signaling including the activity of CNTF (Croker et al., 2008). Additionally, SOCS3 has been shown to negatively regulate interferon (IfN) and other cytokines (Yu et al., 2018). Sun et al., (2011) provided evidence that this pathway is required for the regeneration-promoting effect of SOCS3 in that its effect is lost when Stat3 is also mutated. However, as with Pten, other pathways likely contribute to Socs3-dependent axon regeneration.

### Overcoming type-selective survival and regeneration

RGC types vary dramatically in resilience, with the most resilient and vulnerable types showing ∼99% and ∼1% survival at 14dpc, respectively (Duan et al., 2015: Tran et al., 2019). These molecular differences allowed us to identify targets for neuroprotection (Tran et al., 2019). Here we asked whether neuroprotective and pro-regenerative interventions also selectively affected specific types, which could confer a similar opportunity for target discovery.

In fact, the interventions enhanced survival of most RGC types to similar extents. A few types were rescued with modest selectivity; they included J-RGCs and two types of ooDSGCs. All of these types are inherently vulnerable to ONC (Tran et al., 2019). Conversely, RGCs of the T5 subclass were somewhat poorly rescued. Nonetheless, our overall conclusion is that these interventions showed little selectivity among RGC types, with increased survival scaled to the inherent resilience of each type or subclass.

In contrast, the effects on regeneration varied among interventions. Pten deletion led to selective regeneration of alphaRGCs, consistent with our previous results (Duan et al., 2015), while the additional deletion of Socs3 and overexpression of CNTF was able to overcome these type-specific barriers. This strengthens the hypothesis that broad and extended long-distance regeneration of CNS neurons might only be possible when manipulating multiple genes/factors simultaneously.

### Distinct programs drive death, survival and regeneration

ScRNA-seq revealed transcriptional programs associated with distinct cell states across conditions. In each analysis, three main groups are evident: RGCs that were degenerating or vulnerable, RGCs that survived but failed to extend axons >1mm, and RGCs that not only survived but also extended axons at least 1 mm past the crush site. We identified transcriptional programs associated with each group.

#### Vulnerable and dying RGCs

RGCs in this group expressed genes commonly associated with intrinsic apoptotic signaling and stress response pathways. They included *Ddit3* (CHOP) and *Cepbg*, which are key negative regulators of RGC survival (Tian et al., accompanying paper; Hu et al., 2012; Syc-Mazurek et al., 2017). Most of these genes are globally upregulated by RGCs, including those that are relatively resilient (see also Tran et al 2019).

#### Surviving RGCs

RGCs in this group suppress the degenerative program and upregulate genes implicated in neuronal development, axonogenesis, synaptic organization and synaptic function. Reactivation of developmental genes has been noted in regenerative CNS neurons (Hilton and Bradke, 2017; Poplawski et al., 2020) and synaptic/neuronal activity promotes axon regeneration and functional connectivity following injury as well as suppression of apoptotic signaling pathways (Enes et al., 2010; Hilton et al., 2021; Léveillé et al., 2010; Li et al., 2016; Lim et al., 2016; Tedeschi et al., 2016; Williams et al., 2015; Zhang et al., 2019). We speculate that some of those cells may either initiate a regenerative program that fails or are in an early regenerative state which at the point of collection only enabled short distance regeneration.

#### Regenerating RGCs

These RGCs suppressed injury response and activated axonogenesis-related genes, but additionally activated other pathways, which included previously RAGs (Abe and Cavalli, 2008; Chandran et al., 2016); genes related to the immune response, which have been shown to induce RGC axon regeneration (Benowitz and Popovich, 2011; Bollaerts et al., 2017; Schwartz and Raposo, 2014; Sun et al., 2011), and genes involved in hormone and neuropeptide function. Previous studies have implicated an essential role of hormones in axon growth (Baudet et al., 2009) and neuronal survival (Sanders et al., 2005) during development, and several are known to be upregulated following axotomy of and/to enhance axon outgrowth from peripheral neurons (Holmes et al., 2000; Palkovits, 1995; Tran et al., 2019; Yuan et al., 2010). The appearance of a coordinated neuropeptide-related response in injured CNS neurons, which generally fail to regenerate, along with the gene therapy results (Figure 8, discussed below) is noteworthy from a translational perspective, in that peptides have been used as safe and effective therapeutics in multiple indications.

### Stepwise establishment of responses to injury

By collecting and analyzing cells at several time points, we followed the emergence of gene expression programs regulated by Pten, Socs3 and CNTF. Identifying the dynamics of these programs helps narrow the window of opportunity for therapeutic strategies.

#### Prior to injury

We initiated downregulation of Pten and Socs3 and overexpression of CNTF two weeks before ONC. Although our aim was to ensure that gene expression changes had occurred by the time of injury, this protocol provided an opportunity to distinguish injury-dependent from injury-independent effects. Prior to injury the triple intervention (C/PS_CKO_) led to upregulation of pro-regeneration !RAG” genes previously identified in both peripheral and central nervous systems (Renthal et al., 2020; Yang et al., 2020). These changes mimic the !pre-conditioning” effect seen in DRG injury models (Neumann and Woolf, 1999; Richardson and Issa, 1984). Our data suggests that these changes act in part prior to injury, a measure of little therapeutic value. Despite that caveat, Pten and Socs3 deletion or CNTF overexpression post-injury do elicit potent axon regeneration in some injury models (Danilov and Steward, 2015; Du et al., 2015; Hellström et al., 2011; Sun et al., 2011).

#### Shortly after the injury

At 2dpc, the interventions showed little effect on the injury responses of RGCs (Figure 3H), although SCENIC analysis identified a modest enhancement of regeneration-associated expression patterns compared to those at 0dpc. Thus, even though Pten, Socs3 and CNTF levels have been affected prior to injury, activation of the degeneration program observed in the absence of interventions is not substantially attenuated by the interventions nor are survival programs activated early after injury.

#### One week after the injury

Between 2dpc and 7dpc, all interventions exerted dramatic effects. Pten deletion mitigated the general injury response, by attempting to re-establish the gene expression milieu of uninjured RGCs. RGCs additionally became more distinct in their gene expression profile, with survival and regenerative programs robustly activated. This separation was still apparent at 21dpc, showing that the regenerative state achieved in the first week after the injury is maintained over time.

In short, we demonstrate multiple temporal phases in the effects of intervention. First, there is a modest upregulation of a regenerative state prior to and independent of injury. Second, little change occurs during the first two days after ONC. Third, further degenerative changes are prevented, and survival and regeneration programs are activated between 2 and 7dpc. Finally, expression changes are modest between 7 and 21dpc.

### Promoting axon regeneration

To test if genes correlating with regeneration contribute to this process, we used gain-of-function experiments in which we overexpressed emerging genes present in the regeneration modules identified in this study.

#### Neuropeptides

A key pathway in the !regeneration modules” was annotated as involving hormone and neuropeptide secretion and signaling. We chose two candidates from this group to test - Crh and Gal. Both promoted axon regeneration, as did another peptide, urocortin (Ucn), that we previously identified as selectively expressed by resilient cells (Tran et al., 2019). Further studies will be needed to identify the cellular pathways they affect and to determine whether signaling is cell-autonomous or non-autonomous. Signaling could be direct to RGCs as the Crh receptor (*Crhr1*) is expressed in a subset of them. In contrast, we did not detect the expression of the receptors for Galanin (*GalR1, GalR2* and *GalR3*) in RGCs, suggesting GAL may act cell-non-autonomously or through yet unidentified receptors.

#### Wt1

Wt1 has been shown to exert anti-apoptotic functions in several cell types (Huff, 2011; Loeb, 2006; Yang et al., 2007) including developing RGCs (Wagner et al., 2002). Moreover, Wt1 regulates expression of POU4F2 (Brn-3b), a key regulator of RGC maturation (Wagner et al., 2002) and binds to promoters of multiple genes implicated in axonal regeneration (Gao et al., 2019; Hartl and Schneider, 2019). More recently, studies revealed that Basp1, a growth cone associated protein, is a binding partner of Wt1 (Hartl and Schneider, 2019). In our SCENIC analysis, we noted that genes potentially regulated by and co-expressed with Wt1 include a high proportion of membrane-associated genes implicated in axon outgrowth. Like Pten and Socs3, Wt1 itself is an oncogene and tumor suppressor gene, and therefore problematic as a therapeutic candidate. Notwithstanding, the genes it regulates may provide insights into the control of neuronal survival and axon regeneration as well as a source of novel candidates.

#### Atf3, Atf4, Ddit3 and Cebpg

This set of 4 TFs play key roles in coupling effects of axonal injury to neuronal death; in their absence, neurodegeneration is attenuated (Tian et al., accompanying paper). We observed no effect of deleting any of them on axonal regeneration, supporting the hypothesis that programs regulating survival and regeneration are distinct. Likewise, deletion of *Atf4, Ddit3* or *Cebpg* did not significantly affect the ability of Pten deletion to promote axonal regeneration. On the other hand, Pten deletion was no longer able to promote regeneration when *Atf3* was also deleted. This result supports the idea that Atf3 promotes axonal regeneration of injured peripheral sensory neurons (Renthal et al., 2020) but is seemingly inconsistent with its role in promoting RGC death (Tian et al., accompanying paper). However, its appearance in both degenerative and regenerative modules (MO-M5 and -M6) suggests that unlike the other TFs we assessed it plays a dual role at successive stages of the injury response.

## ACKNOWLEDGEMENTS

We thank Chen Wang, Yiming Zhang and McKinzie E Arnold for assistance and Karthik Shekhar for support in bioinformatic analysis. This work was supported by grants from Wings for Life Spinal Cord Research Foundation to A.J., and NIH grants NS104248-01 to J.R.S., EY029360 to N.T, EY030204-01 to J.R.S and Z.H., EY032181 to F.T. and R01EY021526 and R01EY026939 to Z.H and the Dr. Miriam and Sheldon G. Adelson Medical Research Foundation (to Z.H.) and Gilbert Family Foundation (to Z.H.). Viral cores were supported by the grants from the NIH (HD018655 and P30EY012196).

## AUTHOR CONTRIBUTION

A.J., N.M.T., W.Y., F.T., Z.H and J.R.S. conceived and designed experiments and analyzed data. A.J., N.M.T., I.B., F.T., R.S. performed experiments. Z.H and J.R.S. provided supervision and acquired funding. A.J., N.M.T., W.Y., Z.H. and J.R.S. wrote the paper with input from all authors.

## STAR★METHODS

### KEY RESOURCES TABLE

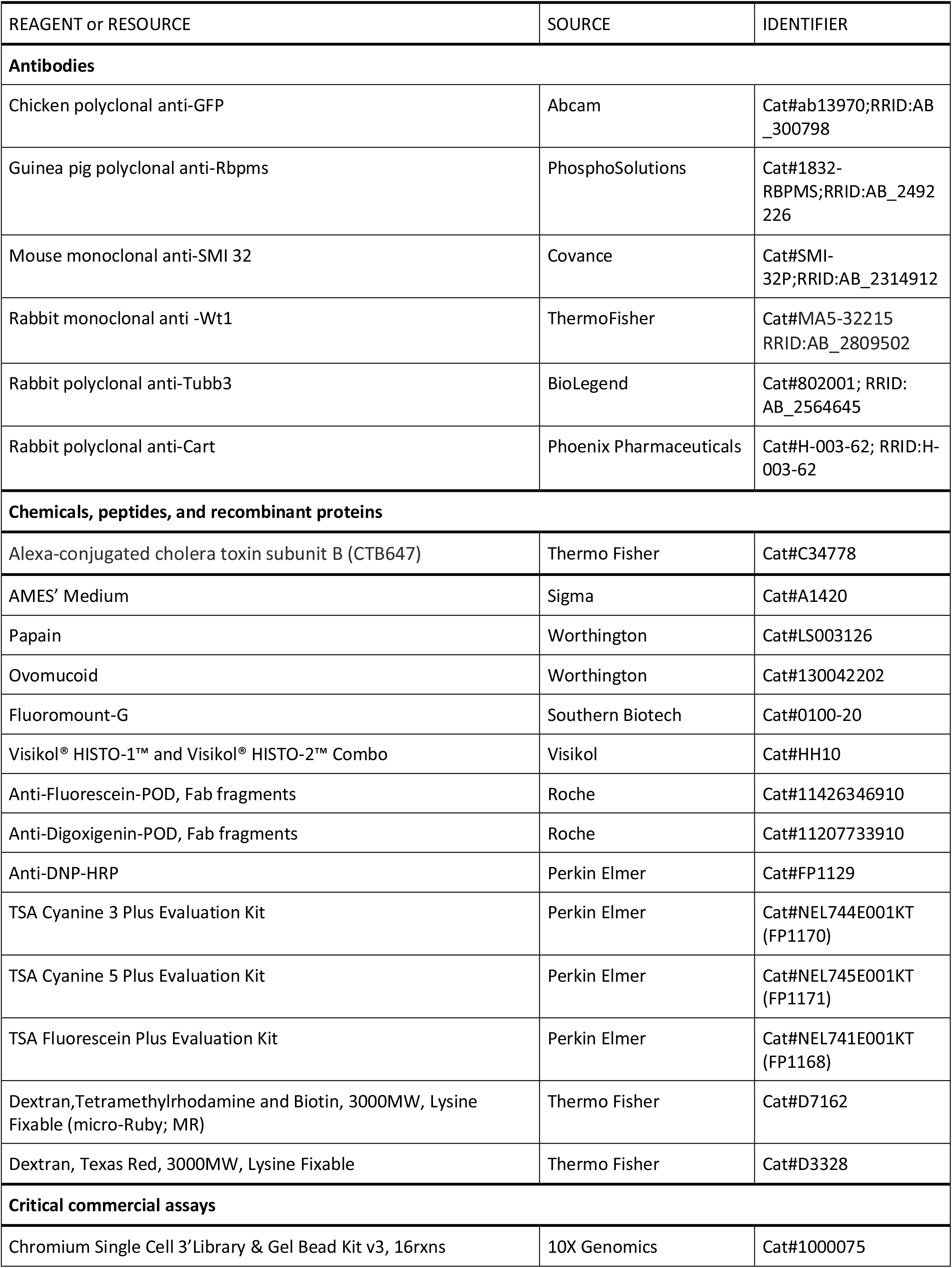
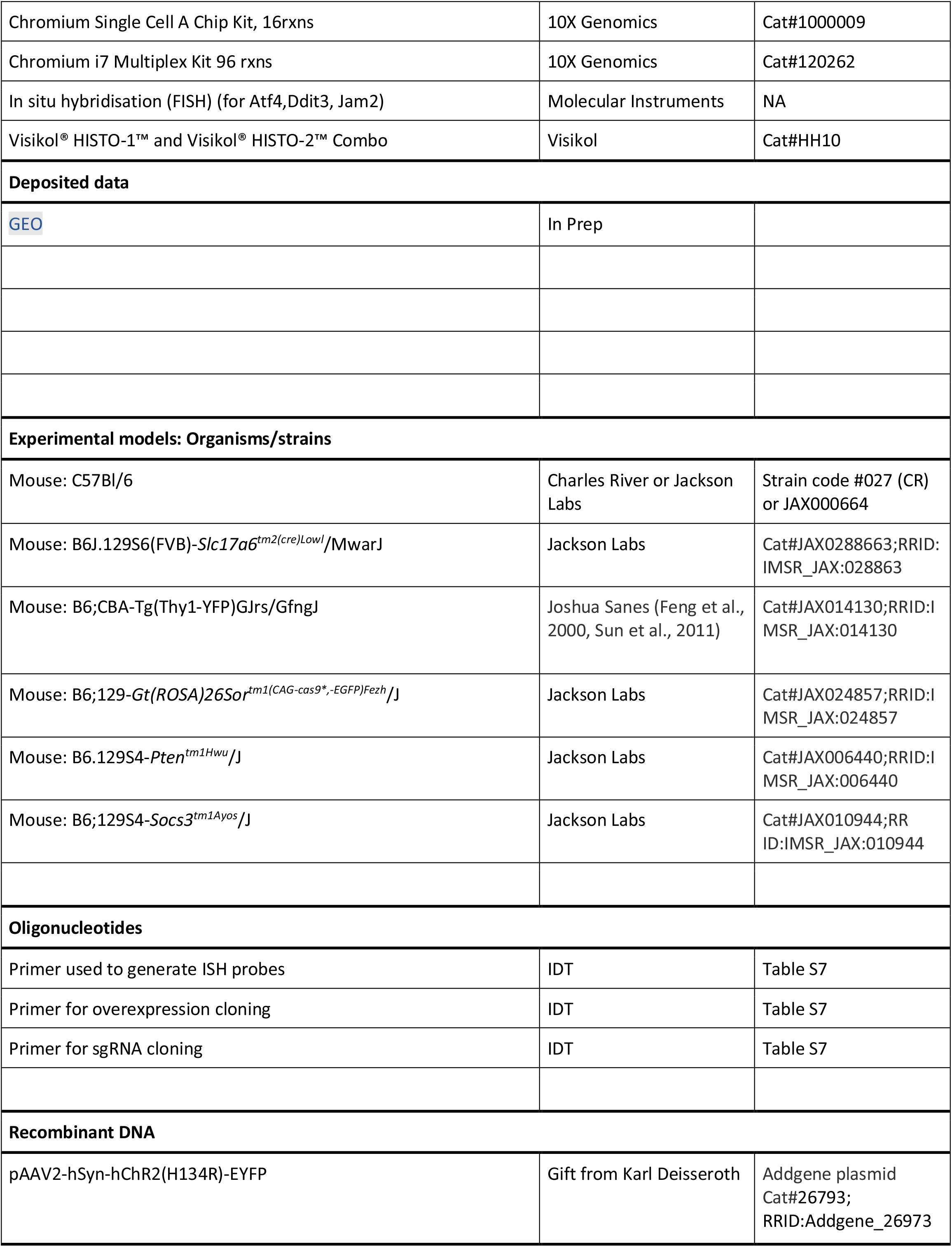
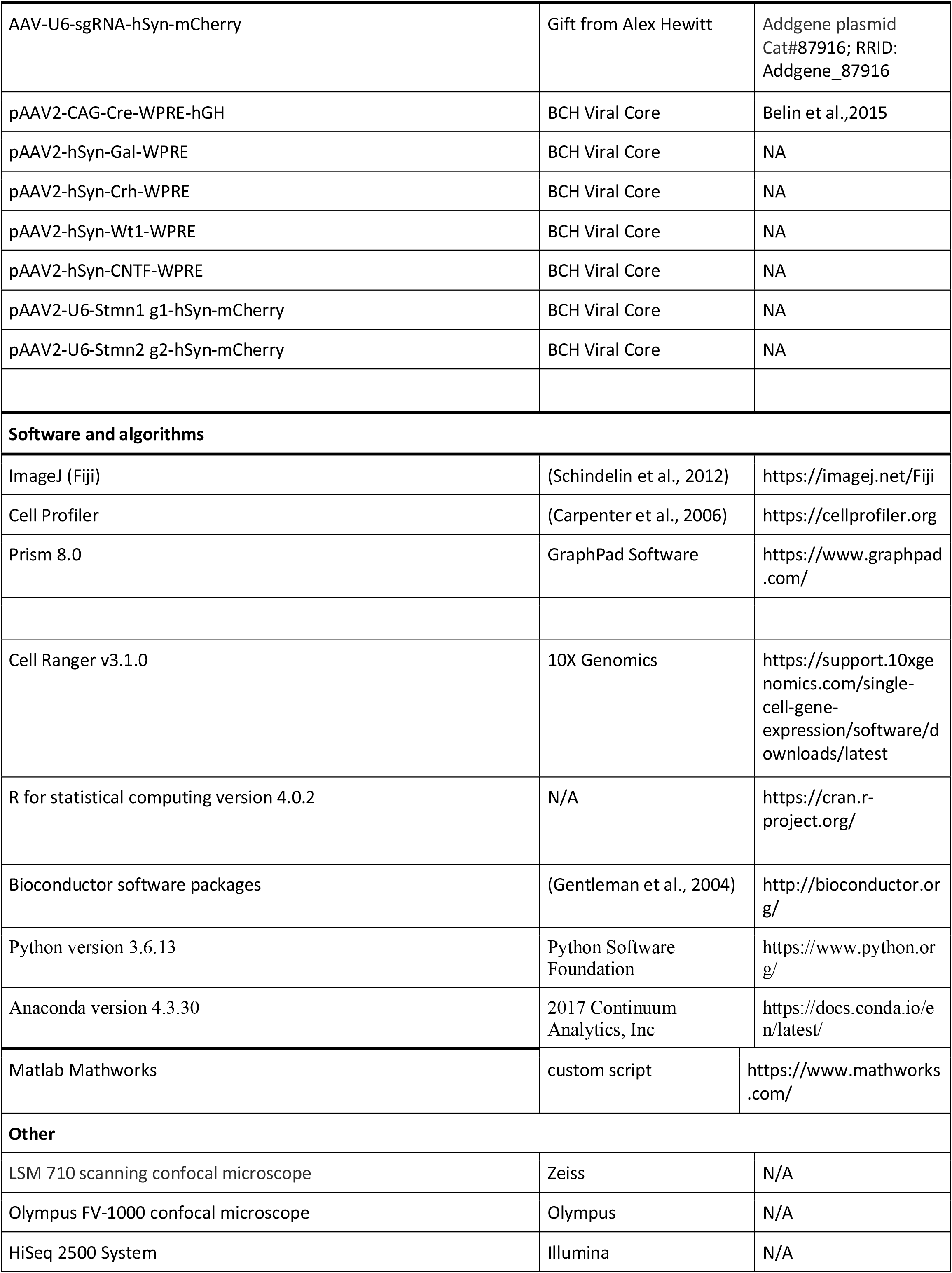
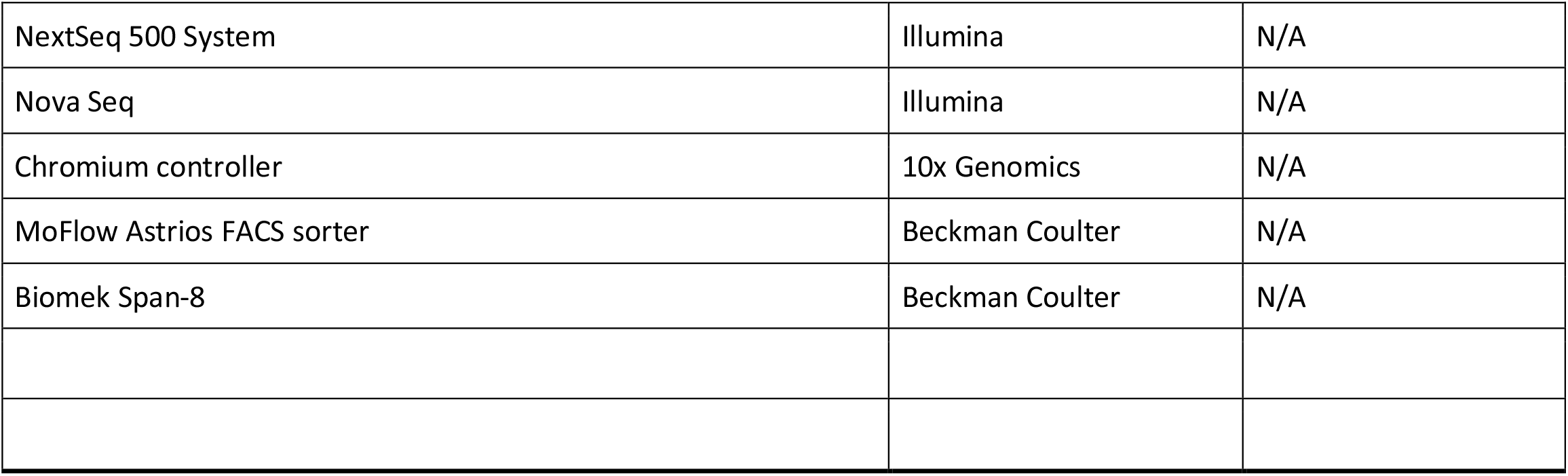

### CONTACT FOR REAGENT AND RESOURCE SHARING

Further information and requests for resources and reagents should be directed to and will be fulfilled by J.R.S.

### EXPERIMENTAL MODEL AND SUBJECT DETAILS

#### Animals

All animal experiments were approved by the Institutional Animal Care and Use Committees (IACUC) at Harvard University and Children’s Hospital, Boston. Male and female mice were used interchangeably. Mice were maintained in pathogen-free facilities under standard housing conditions with continuous access to food and water. All experiments were carried out in adult mice from 4 to 12 weeks of age. The following mouse strains were used:

Pten loxP/loxP (JAX # 006440)

Pten loxP/loxP Socs3 loxP/loxP (Sun et al., 2011)

Thy1-stop-YFP Line #17 (Sun et al., 2011).

Vglut2-Cre (JAX #0288663)

Rosa26-LSL-Cas9 knockin (JAX #024857).

C57Bl/6J (JAX #000664)

### METHOD DETAILS

#### Optic Nerve Crush

Mice were anesthetized with ketamine/xylazine (ketamine 100-120 mg/kg and xylazine 10 mg/kg). We performed optic nerve injury as previously described (Park et al., 2008, Tran et al., 2019). Briefly, the optic nerve was exposed intraorbitally and crushed with fine forceps (Dumont #5 FST) for ∼2s approximately 0.5-1mm behind the optic disc. Eye ointment was applied post-operatively to protect the cornea.

#### Intravitreal injection of AAV

Mice were anaesthetized with ketamine/xylazine (ketamine 100-120mg/kg and xylazine 10 mg/kg) and injected intravitreally with ∼2μl of volume of AAV2 (in 1x PBS) carrying the gene of interest driven by a CAG promoter, or an sgRNA driven by a U6 promoter. Concentration of viruses was adjusted to ∼5 x 10^12. For injections, we first removed ∼2μl intravitreal fluid from the eye with a sterile glass micropipette. Another glass micropipette or a 33-gauge Hamilton syringe was then inserted through the sclera about 0.5 mm posterior to the limbus and into the vitreous chamber without touching the lens. AAV solution (∼2μl) was injected. After injection, antibiotic ophthalmic ointment was applied, and mice were warmed on a heating pad until fully awake.

#### Anterograde tracing of regenerating axons

To assess axon regeneration, axons were anterogradely labeled by intravitreal injection of CTB conjugated with Alexa-647 (Life Technology) 48 hours before sacrifice. After 4% PFA perfusion, heads were post-fixed overnight in 4% PFA. Optic nerves were micro-dissected and meninges surrounding the nerve were removed. Nerves were then cleared using reagents and protocol provided from Visikol®. Briefly, nerves were dehydrated with 100% methanol for 4 minutes and then transferred into Visikol Histo-1 solution for overnight incubation at 4°C. The next day the nerves were incubated in Visikol Histo-2 solution for at least 2 hours before mounting them in Visikol Histo-2 solution and imaged with the LSM710 confocal microscope.

In some cases, we used an iDISCO tissue clearing method (Renier et al., 2014). In this method, optic nerves were incubated in the dark for 0.5h with 80% tetrahydrofuran (THF, Sigma-Aldrich 360589-500ML)/H_2_O and then transferred to 100% THF for 1h. Nerves were then incubated in Dichloromethane (DCM, Sigma-Aldrich 270997-1L) for 20min and switched to dibenzyl ether (DBE, Sigma-Aldrich 33630-250ML) until they were completely transparent (at least 3h).

Optic nerves showing incomplete crush as evidenced by continuous labeling of axons through the chiasm and/or a different morphology then regenerating axons (pearls on a string) were excluded from the analysis; they comprised a low percentage of all nerves analyzed (Tran et al., 2019).

#### Retrograde labeling of regenerating RGCs

Retrograde labeling of regenerated RGCs was performed as described previously (Zhang et al., 2019). Twenty days after ONC, mice were anesthetized and placed in a stereotaxic frame. The crushed optic nerve was exposed using a superior temporal intraorbital approach by drilling a hole into the skull and removing overlying brain tissue. After exposing the optic nerve ∼1.5 mm distal to the crush site, we cut the nerve with a fine blade and delivered 100-300 nL of 5% dextran (3kDa dextran conjugated to biotin; ThermoFisher #D7162 or #D3328) solution diluted in sterile PBS to the stump. We then placed a small piece of gelfoam (Fisher Scientific) soaked in 5% dextran solution on the cut nerve stump. The scalp was sutured, and animals recovered on a heating pad until they regained consciousness. For single cell isolation and SS2, mice were injected with microRuby and perfused ∼24 hours after delivery.

#### Cell preparation and FACS

We used the methods detailed in Tran et al. (2019) for dissociation and FACS sorting of RGCs. Briefly, retinas were dissected in AMES solution (equilibrated with 95% O2/5% CO2), digested in papain, and dissociated to single cell suspensions using manual trituration in ovomucoid solution. For a concentration of 10 million cells per 100μl, 0.5μl of 2μg/μl anti-CD90 (conjugated to various fluorophores) (Thermo Fisher Scientific) was used to stain (15 minutes incubation), washed with an excess of media, spun down and resuspended again in AMES+4%BSA to a concentration of ∼7 million cells per 1 ml. Just prior to FACS the live cell marker calcein blue was added. RGCs were collected based on high CD90, GFP and, in some cases, MicroRuby co-expression. For 10X experiments, cells were collected into ∼100ul of AMES+4%BSA per 25,000 sorted cells. Following collection cells were spun down and resuspended in PBS+0.1% non-acetylated BSA at a concentration range of 500-2000 cells/ul for droplet-based scRNAseq per manufacturer’s instructions (10x Chromium). For SS2 experiments, single cells were collected into 96 well plates filled with 5µl of TCL lysis buffer, containing 1% BME, spun down and frozen at -80°C till further processing.

#### RNA-sequencing

##### 3 ’droplet-based scRNA-seq

Single cell libraries were prepared using the Single-cell gene expression v2/v3 kit on the Chromium platform (10X Genomics, Pleasanton, CA) following the manufacturer’s protocol. Briefly, single cells were partitioned into Gel beads in EMulsion (GEMs) in the Chromium instrument followed by cell lysis and barcoded reverse transcription of RNA, amplification, enzymatic fragmentation, 5 ‘adaptor attachment and sample indexing. On average, approximately 8,000-12,000 single cells were loaded on each channel and approximately 3,000-7,000 cells were recovered. Libraries were sequenced on Illumina HiSeq 2500, or NovaSeq platforms (Paired end reads: Read 1, 26 bases, Read 2, 98 bases).

#### Retrograde labeled RGCs: Smart-seq2

We generated RNA-Seq libraries using a modified Smart-seq2 method (Picelli et al., 2014) with the following minor changes: Before running RT, RNA was purified using 2.2X SPRI-beads (Beckman Coulter, A3987) followed by 3 wash steps with 80% EtOH, elution in 4µl of RT primer mix and denatured at 72°C for 3 min. Six microliters of the first-strand reaction mix, containing 0.1μl SuperScript II reverse transcriptase (200 U/μl, Invitrogen), 0.25μl RNAse inhibitor (40 U/μl, Clontech), 2μl Superscript II First-Strand Buffer (5x, Invitrogen), 0.1μl MgCl_2_ (100 mM, Sigma), 0.1μl TSO (100 μM) and 3.45μl Trehalose (1M), were added to each sample. Reverse transcription reaction was carried out by incubating at 50°C for 90 min and inactivation by incubation at 85°C for 5 min. After PCR preamplification, PCR was purified using 0.8X of AMPure XP beads (Beckman Coulter), with the final elution in 12μl of EB solution (Qiagen). For tagmentation the Nextera DNA Sample Preparation kit (FC-131-1096, Illumina) was used, and final PCR was performed as follows: 72°C 3 min, 95 °C 30 s, then 12 cycles of (95°C 10 s, 55°C 30s, 72°C 1min), 72°C 5min. Purification was done with a 0.9X of AMPure XP beads. Libraries were diluted to a final concentration of 2nM, pooled and sequenced on Next-Seq 500 or Nova-Seq, 50bp paired end.

#### Whole mounts

Eyes were either collected from animals intracardially perfused with 15-50ml of 4% paraformaldehyde (PFA), and post-fixed for an additional 15 minutes, or dissected from nonperfused animals and immersion fixed in 4% PFA for 30 minutes. Eyes were transferred to PBS until retinas were dissected.

To immunostain whole mounts, retinas were incubated in blocking solution (5% normal serum, 0.3% Triton-X100 in PBS) for 3 hours, followed by incubation with primary antibodies (in blocking solution) for 5-7 days, and secondary antibodies (in PBS) overnight. All incubations were done at 4°C with gentle rocking.

*Jam2* expression was assessed by fluorescent *in situ* hybridization using a hybridization chain reaction method (https://www.moleculartechnologies.org/). Probes of 15-20nt length were generated by Molecular Instruments. Retinas were dissected in RNAse-free 1xPBS and immediately washed 2 x 5min in PBST (1xPBS + 0.1% Tween20) on ice. Retinas were then dehydrated using MeOH – PBST mix series (0%, 25%, 50%, 75% and 100% of MeOH), each step for 15min on ice. Retinas were incubated in 100% methanol overnight at -20°C. After rehydration on ice the next day (inverted order of previous dehydration) and 10min incubation in PBST at RT, retinas were incubated for 30min at 37°C for pre-hybridization. Then retinas were incubated in hybridization buffer including the probe (2.5nM) overnight at 37°C. After hybridization retinas were washed 4 x 15min with wash buffer (at 37°C) followed by 2x 5min in 5x saline-sodium citrate (SSC) at room temperature (RT). The amplification step was performed with amplifiers B1 or B2for 24hrs at RT in the dark. Finally, retinas were immunostained for RBPMS as above and mounted.

#### Retinal sections

Eyes were collected and retinas dissected as described above. Retinas were then sunk in 30% sucrose, embedded in tissue freezing media, and cryosectioned at 20μm. For IHC, slides were incubated for 1 hour in protein block, primary antibody incubation overnight, and secondary antibodies for 2-3 hours. Initial block and secondary antibody incubation were done at RT and primary antibody incubation at 4°C.

For FISH, probes were either obtained from Molecular Instruments (*Atf4*, *Ddit3* and *Jam2*) and used as described previously (Li et al., 2020). All other probes were generated, and FISH was performed as described in Tran et al. (2019).

#### Design of overexpression and knockdown vectors

Vectors were cloned by Synbio Technologies (Monmouth Junction, NJ 08852) using the pAAV-hSyn-hChR2(H134R)-EYFP plasmid (Addgene #26973) to replace the hChrR2(H134R)-EYFP with the gene target sequence for over expression experiments. Virus of serotype AAV2/2 was then generated by the Boston Children’s Hospital Viral Core. For Crispr mediated KD a modified AAV-U6-sgRNA-hSyn-mCherry plasmid (Addgene #87916) was used. The AAV2-based Crispr/Cas9 approach we employ here has been established as an effective modality for somatic knockdown in adult mouse RGCs (Hung et al., 2016). To account for possible off target effects, we delivered a mix of 5 AAV2 single-guide RNA (sgRNA) expression vectors to the eyes of mice that express Cas9 specifically in RGCs (VGlut2-Cre; LSLCas9-eGFP), which lead to high infection rates as described previously and indicated in Figure S7A. Vectors and sequences used for manipulation experiments are displayed in the Key Resources Table.

#### Computational Methods

##### Reads alignment of 3 ’droplet-based scRNA-seq data

Sequenced reads were demultiplexed using cellranger (version 2.1.0, 10x Genomics) !mkfastq” function and aligned to mouse genome mm38 with modified transcriptome (Tran et al., 2019) using cellranger !count” function.

##### Clustering and cell type identification in 3 ’droplet-based scRNA-seq data

The generated gene count matrix was processed using the R package !Seurat” (Version 4.0.1). Both the standardized log normalization and the “sctransform” method were used for processing. Briefly, the gene expression matrix generated by log normalization and scaling with a factor of 10000 were used for differential gene expression analysis and data visualization, while the sources of variation for each intervention at each time point were removed using the “sctransform” framework and clustering was performed based on the corrected values. The 2000 top ranked common features among interventions at each time point were selected using function “SelectIntegrationFeatures”. The canonical correlation analysis-based data Integration method (function “IntegrateData”) was applied using each and all the interventions without optic nerve crush as the reference dataset. Principal Component analysis (PCA) was performed, and the top 100 PCs were used to construct a shared nearest neighbor (SNN) graph, with k=100 as the k-nearest neighbors. The Louvain algorithm with multilevel refinement algorithm was used for modularity optimization in identifying the clusters. In the first round of clustering, canonical retinal cell class markers were used to identify major cell classes in the dataset including amacrine cells and RGCs. Only RGCs were retained for further analysis, using the pipeline described above. Results were evaluated based on number of distinct marker genes in each cluster as well as their correspondence to the RGC type prediction using machine learning algorithm XGBoost (See below for details). A resolution of 3 was chosen as the clustering parameter in the function “FindClusters” based on the integrated SNN.

To identify RGC types in each cluster, two methods were used. First, a machine learning algorithm “XGBoost” was applied as described previously (Yan et al., 2020; van Zyl et al., 2020) to build the RGC type predictor based on the RGC atlas dataset (Tran et al., 2019). Confusion matrices were generated between the predicted result and clusterings at various resolutions. High consensus was observed among results, with subtle differences in a small set of clusters. In those cases, differential gene expression (DGE) analysis was performed to verify the sub-division of certain types using the !MAST” method by !Seurat” function !FindMarkers” based on the log normalized data. Clusters were kept as separate if more than 5 differentially expressed (DE) genes were identified in both groups with over 10% expression in either cluster and over 0.5 log-fold change. Otherwise, the clusters under evaluation were merged or a lower resolution was chosen. As a second measure to ensure the accuracy of RGC type identification, expression of type marker genes from (Tran et al., 2019) was inspected in each cluster. When measuring at the level of subclasses, the composition of RGC types was defined as in Figure 2J in Tran et al., 2019.

##### Detecting of Cre/CNTF transcript expression in dataset

A separated alignment of the reads was performed using the same cellranger count function but using a reference genome to which the WPRE (Cre-HAtag-WPRE) sequence, shared by both AAV vectors (Cre and Cntf) had been added.

Cre-HAtag-WPRE sequence:

tccaatttactgaccgtacaccaaaatttgcctgcattaccggtcgatgcaacgagtgatgaggttcgcaagaacctgatggacatgttcagggatcgccaggcgttttctgagcatacctggaaaatgcttctgtccgtttgccggtcgtgggcggcatggtgcaagttgaataaccggaaatggtttcccgcagaacctgaagatgttcgcgattatcttctatatcttcaggcgcgcggtctggcagtaaaaactatccagcaacatttgggccagctaaacatgcttcatcgtcggtccgggctgccacgaccaagtgacagcaatgctgtttcactggttatgcggcggatccgaaaagaaaacgttgatgccggtgaacgtgcaaaacaggctctagcgttcgaacgcactgatttcgaccaggttcgttcactcatggaaaatagcgatcgctgccaggatatacgtaatctggcatttctggggattgcttataacaccctgttacgtatagccgaaattgccaggatcagggttaaagatatctcacgtactgacggtgggagaatgttaatccatattggcagaacgaaaacgctggttagcaccgcaggtgtagagaaggcacttagcctgggggtaactaaactggtcgagcgatggatttccgtctctggtgtagctgatgatccgaataactacctgttttgccgggtcagaaaaaatggtgttgccgcgccatctgccaccagccagctatcaactcgcgccctggaagggatttttgaagcaactcatcgattgatttacggcgctaaggatgactctggtcagagatacctggcctggtctggacacagtgcccgtgtcggagccgcgcgagatatggcccgcgctggagtttcaataccggagatcatgcaagctggtggctggaccaatgtaaatattgtcatgaactatatccgtaacctggatagtgaaacaggggcaatggtgcgcctgctggaagatggcgattacccatacgatgttccagattacgcttaaTCTAGAGTCGACCTGCAGAAGCTTatcgaTaatcaacctctggattacaaaatttgtgaaagattgactggtattcttaactatgttgctccttttacgctatgtggatacgctgctttaatgcctttgtatcatgctattgcttcccgtatggctttcattttctcctccttgtataaatcctggttgctgtctctttatgaggagttgtggcccgttgtcaggcaacgtggcgtggtgtgcactgtgtttgctgacgcaacccccactggttggggcattgccaccacctgtcagctcctttccgggactttcgctttccccctccctattgccacggcggaactcatcgccgcctgccttgcccgctgctggacaggggctcggctgttgggcactgacaattccgtggtgttgtcggggaagctgacgtcctttccatggctgctcgcctgtgttgccacctggattctgcgcgggacgtccttctgctacgtcccttcggccctcaatccagcggaccttccttcccgcggcctgctgccggctctgcggcctcttccgcgtcttcgccttcgccctcagacgagtcggatctccctttgggccgcctccccgc

##### Reads alignment and analysis of plate-based full-length Smart-Seq2 dataset

Raw reads were first trimmed by Trimmomatic (version 0.39) and then aligned to GRCm38 (Genome Reference Consortium Mouse Build 38) downloaded from (https://cloud.biohpc.swmed.edu/index.php/s/grcm38_tran/download) using Hisat2 (version 2.1.0). Gene expression matrices for each cell were quantified using featureCounts with GRCm38 transcriptome file version 81. Low quality cells were filtered out using the following criteria: >= 500,000 total reads mapped to genome, >= 1500 genes detected in each cell, >= 40% of reads mapped to the transcriptome. Count matrix calculated with Reads Per Kilobase of transcript, per Million mapped reads (RPKM) was generated for all the cells passed the filter. A similar analysis pipeline was applied for downstream analysis. To identify the RGC type to which each cell belonged, two methods were used. First, a type predictor was built from the droplet-based dataset (Tran et al., 2019) using the “XGBoost” algorithm. Second, correlation analysis was performed to seek the most similar type for each cell based on the overall gene expression. The two methods yielded consistent results, indicating reliable RGC type identification. To quantify regenerating RGC subclass contribution, numbers arising from the correlation analysis were used.

##### Measure expression of gene sets

The overall expression of gene sets identified previously (Tran et al., 2019) or from current study were measured using gene set scores. First, genes in the list were filtered to remove those in < 25% of cells of all types. Then, for each cell, the mean expression value of the genes in the set j (Exp_i,j_) and in the total transcripts (Exp_i_) were calculated. Next, the score of gene set j in cell i (S_i,j_) was calculated as = Exp_i,j_ - Exp_i_. Finally, the averaged score of gene set j in each group was visualized in Figure 3B,H and Figure S4I.

#### Co-expression gene modules

We used Monocle3 (Cao et al., 2019; Qiu et al., 2017; Trapnell et al., 2014) to examine gene co-expression modules in our scRNA-seq dataset. The dataset was pre-processed using the “preprocess_cds ’function (num_dim = 100) by the “PCA” method and dimensionality was reduced using the $reduce_dimensions “function. Batch differences were corrected by the MNN method using the ‘align_cds ’(dimensions = 100) function and dimension reduction was repeated (Haghverdi et al., 2018). Cells were clustered using the !louvain” method (Levine et al., 2015) using the $cluster_cells ’function (k=100) and plotted by ‘UMAP’. Cluster-specific genes were identified using the ‘top_markers ’function. To find gene co-expression modules, we input the resulting cluster-specific genes and used the $find_gene_modules ‘(resolution = 1e-2). This resulted in 6 gene expression modules (Table S5), which we then plotted for single cells using ‘UMAP ’ or by cluster using ‘pheatmap’.

##### Transcriptional regulatory network analysis using Scenic

To identify transcriptional regulatory networks, we applied the computational method “Scenic” (Aibar et al., 2017; Van de Sande et al., 2020) to cells collected at each time point separately using expression matrix of all the genes. The function !SCENICprotocol” was run with Nextflow using the singularity image. The list of TFs, genome ranking databases and motif to transcription factor annotations database for mm10 were downloaded from https://resources.aertslab.org/cistarget/. For gene ranks, the 10kb upstream and 10k downstream around the transcription start site were used as search space. Visualization of the result was performed using customized R and Python scripts.

#### GO-pathway analysis of gene lists

Gene ontology analysis was performed on the DE gene lists generated by group comparisons. Cut offs for inclusion of genes obtained from DE analysis are indicated in the individual tables (S2-S5). Ensemble based annotation package “EnsDb.Mmusculus.v79” and Genome wide annotation for Mouse “org.Mm.eg.db” were used by R package “clusterProfiler” (Yu et al., 2012) to identify the enriched pathways.

### QUANTIFICATION AND STATISTICAL ANALYSIS

#### Retinal whole mounts

RGC density was quantified by immunostaining retina whole mounts with an antibody against RBPMS, a pan RGC marker (Rodriguez et al., 2014). Retinas were examined with epiflourescent illumination. Each quadrant was checked for signs of injection site damage, inflammation, or other damage and areas with obvious damage or inflammation were excluded from further analysis. For imaging, the temporal quadrant was avoided because it has a high density of ɑ alphaRGCs (Bleckert et al., 2014), which, as described in results, are resilient to injury. The entirety of one of the other three quadrants was imaged by a tiled Z-stack scan by confocal microscopy on either a Zeiss 710 or Olympus Fluoview1000 scanning laser confocal microscope. A maximum projection spanning the ganglion cell layer was obtained, and the image background was adjusted using the ’normalize local contrast ‘filter in ImageJ. Quantification of RBPMS density used a semi-automated counting method, as previously described in Tran et al. (2019). Briefly, the processed image was thresholded by the $Otsu ’method using Cell Profiler 4.0 (Carpenter et al., 2006) to identify regions-of-interest (ROIs) that demarcated RBPMS+ cells. The resulting ROIs were then exported to a TIFF file and the centroid position of each ROI was determined using the $analyze particles ’function in ImageJ and an overlay of the original image, ROI outline, and centroid position was produced. These overlayed images were further analyzed in Matlab to determine the density of RGCs in each quadrant using custom scripts. For each retinal quadrant, the bounding regions of the quadrant were interactively selected by the user, avoiding the quadrant edges, which can have increased autofluorescence, and areas with minor damage from dissection. Mean densities were calculated as RGCs/mm^2^. Significance was determined by Student’s t-test and p-values were FDR adjusted for multiple comparisons.

#### Retinal sections (CARTPT vs. Gal)

The fluorescent intensity of CART immunostaining and Gal in situ hybridization probe staining were quantified as previously described (Carpenter et al., 2006; Yan et al., 2020). Briefly, three stained sagittal retinal sections were imaged for each condition by confocal microscopy (Zeiss 710) at 40x magnification. Z-stacked images were analyzed in ImageJ. Custom ImageJ macros were used to place circular regions of interest (ROIs; diameter, 3.44 µm) over all cell nuclei/somas that were positive for at least one marker. Fluorescent probe intensity was measured for each marker in each ROI in single Z-slice images. Values were background subtracted using a collection of ROIs negative for both markers. Correlation co-efficient values were calculated in Microsoft Excel and compared to randomized data.

#### Axon regeneration

The cleared, whole nerve was imaged with a 20X air objective, zoom 1x. From the center of the nerve, 7 single stacks (2μm stack size) were maximum projected to a total volume of 14μm per nerve. After defining the crush site, lines spaced equidistant from each other at 500μm intervals from the crush site to where the longest axon could be detected were introduced for bin-by-bin axon quantification. As described previously (Duan et al., 2015; Park et al., 2008), we quantified the total number of regenerating axons, Σad, using the formula Σad = πr2x [average axons/mm]/t, where the total number of axons extending distance d in a nerve having a radius of r was estimated by summing over all sections with thickness t. Bin-by-bin axon quantification was used to assess significant differences between individual distances (500, 1000, 1500, 2000μm) was determined by two-way ANOVA or mixed effects analysis followed by Dunnett’s multiple comparison test with GraphPad Prism, p < 0.05 = *.

For CRISPR-Cas9 mediated KO candidates, maximum projections of Z-stack images were used to capture all regenerated axons. For image analysis, fluorescent intensity profiles along the nerve were generated by the built-in function of ImageJ: Analyze/Plot Profile. To calculate the integral of fluorescent intensity across the entire length of the nerve, a Matlab algorithm was developed (Tian et al., XXX) to quantify the !area under curve” from the plot profile data generated by ImageJ.

### DATA AND SOFTWARE AVAILABILITY

Submission of all the raw and processed datasets reported in this study has been initiated to the Gene Expression Omnibus (GEO) with accession number in process.

### SUPPLEMENTARY DATA

Table S1_Datasets used in this study

Table S2_ DE genes from interventions prior to nerve crush

Table S3_ DE genes from comparison of microRuby positive and negative RGCs

Table S4_DE genes at 7dpc from Pseudo-bulk analysis

Table S5_DE genes at 7dpc from Monocle Modules

Table S6_Regulons (TFs and their regulated genes) at 7dpc from Scenic analysis

Table S7_Oligonucleotides for in situ and virus production

**Figure S1: Manipulation and survival of injured RGCs**

**A)** Percentage of RGCs in which we detected AAV-derived transcripts in indicated datasets.

**B)** Knock out of Pten RNA in uninjured P_CKO_ retina. Pten detected by *in situ* hybridization; all RGCs stained with anti-RBPMS. Scale bar = 50µm.

**C)** Dotplot showing expression of RGC type-specific marker genes (columns) at 7dpc (rows; see color bars, top and right). The size of each circle is proportional to the percentage of cells expressing the gene, and the color depicts the average normalized transcript count in expressing cells.

**D)** Proportion of three RGC types (12_NT ooDS, and 14, CCK-expressing_ooDS, and J-RGCs) at 0 and 7dpc as determined by scRNA-seq. Frequency of both ooDSGC types was increased by all three interventions, but J-RGC frequency increased only in P_CKO_ retina.

**E)** Jam2 expression determined by in situ hybridization in retinal whole mounts at 7dpc (LEFT). Quantification shows significantly increased frequency in P_CKO_ (Mean ’SD, Student#s t-test p < 0.018). Scale bar = 100µm.

**F-H)** Scatterplots showing relative frequencies of RGC subclasses in P_CKO,_ C/P_CKO,_ and C/PS_CKO_ retinas compared to wildtype at 7dpc. (r2 = R_Pearso_ ^2^ values).

**Figure S2: Retrograde labeling and isolation of regenerating RGCs after crush**

**A)** Retrograde labeling of regenerating RGCs. Injection of a 5% dextran into an intact (WT – 0dpc) optic nerve leads to ∼95% high labeling efficiency of RGCs. No labeling was seen in the WT injected control after crush (WT – 14dpc). Regenerating axons are labeled in P_CKO_ retina at 14dpc. Arrows indicate co-labeled RGCs. Green: RBPMS; Red: 5% Dextran; Blue: DAPI.

**A’)** Quantification of retrogradely labeled RGCs from images such as those in **A**. Error bars: SD, n = 3-5. Scale bar = 50µm.

**B)** FACS plots showing gates used to isolate regenerating MR-labeled RGCs. YFP (all RGCs) and MR-fluorescence (regenerating RGCs) were measured. LEFT panel: high number of MR^+^ RGCs after retrograde labeling in non-crushed optic nerve. MIDDLE panel: No MR^+^ detected after injection into a non-regenerating crush condition. RIGHT panel: MR^+^ RGCs after P_CKO_ induced regeneration. Outlines show representative gates used to collect of MR+ and MR-RGCs.

**Figure S3: I*n vivo* validation of injury independent upregulation of selected genes**

**A)** Immunohistochemistry of TUBB3 in retinal cross-sections shows upregulation in C/PS_CKO_ retinas at 0dpc. Scale bar = 50µm.

**B)** In situ hybridization of Stmn1 expression in retinal cross-sections high expression in WT compared to C/PS_CKO_ at 0dpc and 7dpc. Scale bar = 100µm (0dpc), 50µm (7dpc).

**C)** ViolinPlot of Stmn1 expression in C/PS_CKO_ RGCs at indicated times.

**Figure S4: Pseudo-Bulk analysis module GO pathways**

**A)** Results of hypergeometric tests indicating relationships of modules derived from intervention-specific and incremental comparisons among interventions. Statistically significant p-values are shown.

**B)** In situ hybridization with probes for Atf4 and Ddit3 in cross-sections of wildtype or C/PS_CKO_ retina at 7dpc. green/yellow: RGCs in RGC layer, blue: DAPI, red/purple: Ddit3/Atf4. Scale = 50µm.

**C,F)** Top10 GO-pathways enriched in PB-M2 (**C**) and PB-M3 (**F**), performed on intervention specific DE genes (logFC > 0.3, FDR < 0.001)

**D,G)** Cnet plot of Top10 pathways for PB-M2 (**D**) and PB-M3 (**G**) with associated genes. Color of dots represent fold change of genes. Size of the grey dots refer to the number of genes enriched with the GO-term.

**E)** Dotplot showing expression of selected neurotransmitter release genes from PB-M2 at 7dpc.

**H)** Dotplot showing expression of selected cytoplasmic translation genes from PB-M3 at 7dpc.

**I)** Regeneration scores, as in Figure 3B, compiled from 306 genes selectively expressed in MR+ RGCs in C/PS_CKO_.

**J)** Immunohistochemistry of TUBB3 expression in retinal cross-sections in wildtype or C/PS_CKO_ at 7dpc. yellow: RGCs in RGC layer, purple: Tubb3. Scale = 50µm.

**Figure S5: Modules derived from Monocle analysis**

**A)** UMAP distribution of Monocle clustering at 7dpc by intervention.

**B)** In situ hybridization for Gal and Cartpt showing increased expression at 7dpc in C/PS_CKO_ compared to WT. Scale bar = 50µm.

**C)** Levels of Gal and Cartpt expression in C/PS_CKO_ RGCs showing substantial segregation into two populations (correlation

coefficient = -0.5). Scale bar = 25µm.

**D - I)** Cnet plots of Top10 pathways for each Monocle module with associated genes. Color of dots represent fold change of genes.

**Figure S6: Characterization of RGCs that did not map to RGC types**

**A)** Proportions of RGCs in Seurat clusters A-G (see Figure 1D) at 7dpc derived from the indicated datasets.

**B, C)** Dotplots showing enriched expression of selected cell death associated genes in Seurat cluster B (**B**) and selected RAGs genes in Seurat cluster A at 7dpc compared to all other clusters 7dpc.

D) Results of a hypergeometric test showing correspondence of genes in modules derived from Monocle and those in Seurat clusters A-G. Statistically significant p-values are shown.

**E, F)** Top10 GO-pathways enriched in clusters D and F at 7dpc.

**G)** Top20 Regulons based on Regulon Specificity Score (RSS) for each intervention at 7dpc.

**H, I)** Heatmap of key transcription factor (TF) expression and their putative downstream targets in each cell at 0dpc (**H**) and 2dpc (**I**) established by SCENIC analysis. Each row is a single RGC, with color bar at left indicating intervention type. Each column is a single regulon, with key TF listed at bottom. Dotted lines indicate modules discussed in the text.

**Figure S7: Effects of selected genes on RCG axon regeneration**

**A)** Expression of Crh or Gal at 21dpc following AAV infection. Green = RGCs, Red/grey = target gene, Blue = DAPI. Scale bar = 50µm.

**B)** UMAP of Atf3 expression at 7dpc.

**C)** Maximum projections of cleared optic nerves showing anterograde-labeled RGC axons at 12dpc following CRISPR mediated KO of Atf3, Atf4, Ddit3 and Cebpg. Scale bar = 250µm, red cross indicates crush site.

**D)** Quantification of axon regeneration from images such as those in **C**. Deletion of these genes fails to promote regeneration. Data are shown as Mean %SEM, n = 4-6.

**E)** Anterograde-labeled RGC axons following knockdown in P_CKO_ retinas. Data are shown as mean ’SEM. Scale bar = 250µm, red cross indicates crush site.

**F)** Quantification of axon regeneration of P_CKO_ combined with indicated treatment. adjusted p-value: * < 0.05, ** < 0.01, *** < 0.001. 2-way ANOVA or mixed effects analysis with Dunnett’s correction.

**G)** Total RGC density (RBPMS+/mm^2^ cells; mean %SD) in whole mounts following interventions in P_CKO_ shown in F. n=20 in total; *adjusted p-value <.05 (FDR).

**H)** IHC in retinal whole mounts for RBPMS shows increased survival of RGCs at 21dpc following OE-Gal, OE-Wt1 and OE-Crh in P_CKO_. Scale bar = 100µm.

**I)** In situ hybridization of *Cartpt* (MO-M2) in retrogradely labeled tissue from C/PS_CKO_ shows no co-expression with regenerating RGCs. In contrast, genes associated with regeneration (Gal, Wt1 and Crh; MO-M5) show strong expression in regenerating RGCs at 21dpc. Green = RGCs, Red = MR, grey = target gene. Scale bar = 50µm.

