## Supplementary Figures 1-7 for "Overlapping transcriptional programs promote survival and axonal regeneration of injured retinal ganglion cells"

[6]Corresponding authors

[7]Lead contact

#### SUPPLEMENTARY FIGURE LEGENDS

##### Figure S1: Manipulation and survival of injured RGCs

**A)** Percentage of RGCs in which we detected AAV-derived transcripts in indicated datasets.

#### Figure S7: Effects of selected genes on RCG axon regeneration

**A)** Expression of Crh or Gal at 21dpc following AAV infection. Green = RGCs, Red/grey = target gene, Blue = DAPI. Scale bar = 50µm.
**B)** UMAP of Atf3 expression at 7dpc.
**C)** Maximum projections of cleared optic nerves showing anterograde-labeled RGC axons at 12dpc following CRISPR mediated KO of Atf3, Atf4, Ddit3 and Cebpg. Scale bar = 250µm, red cross indicates crush site.
**D)** Quantification of axon regeneration from images such as those in **C**. Deletion of these genes fails to promote regeneration. Data are shown as Mean ±SEM, n = 4-6.
**E)** Anterograde-labeled RGC axons following knockdown in $P_{CKO}$ retinas. Data are shown as mean ±SEM. Scale bar = 250µm, red cross indicates crush site.
**F)** Quantification of axon regeneration of $P_{CKO}$ combined with indicated treatment. adjusted p-value: * < 0.05, ** < 0.01, *** < 0.001. 2-way ANOVA or mixed effects analysis with Dunnett's correction.
**G)** Total RGC density (RBPMS+/$mm^2$ cells; mean ±SD) in whole mounts following interventions in $P_{CKO}$ shown in F. n=20 in total; *adjusted p-value <.05 (FDR).
**H)** IHC in retinal whole mounts for RBPMS shows increased survival of RGCs at 21dpc following OE-Gal, OE-Wt1 and OE-Crh in $P_{CKO}$. Scale bar = 100µm.
**I**) In situ hybridization of *Cartpt* (MO-M2) in retrogradely labeled tissue from $C/PS_{CKO}$ shows no co-expression with regenerating RGCs. In contrast, genes associated with regeneration (Gal, Wt1 and Crh*;* MO-M5) show strong expression in regenerating RGCs at 21dpc. Green = RGCs, Red = MR, grey = target gene. Scale bar = 50µm.

### Suppl.Figure 1

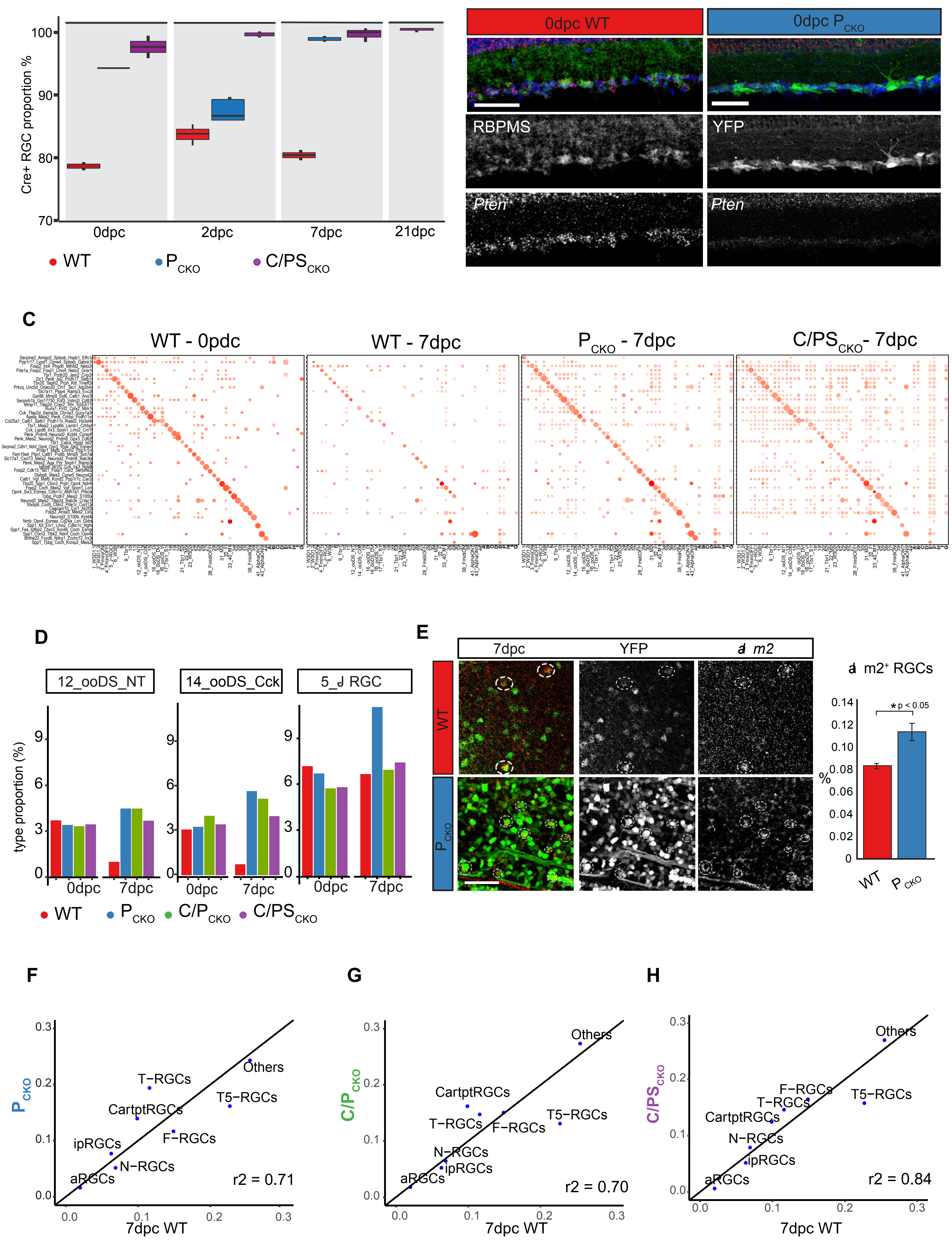

### Suppl. Figure 2

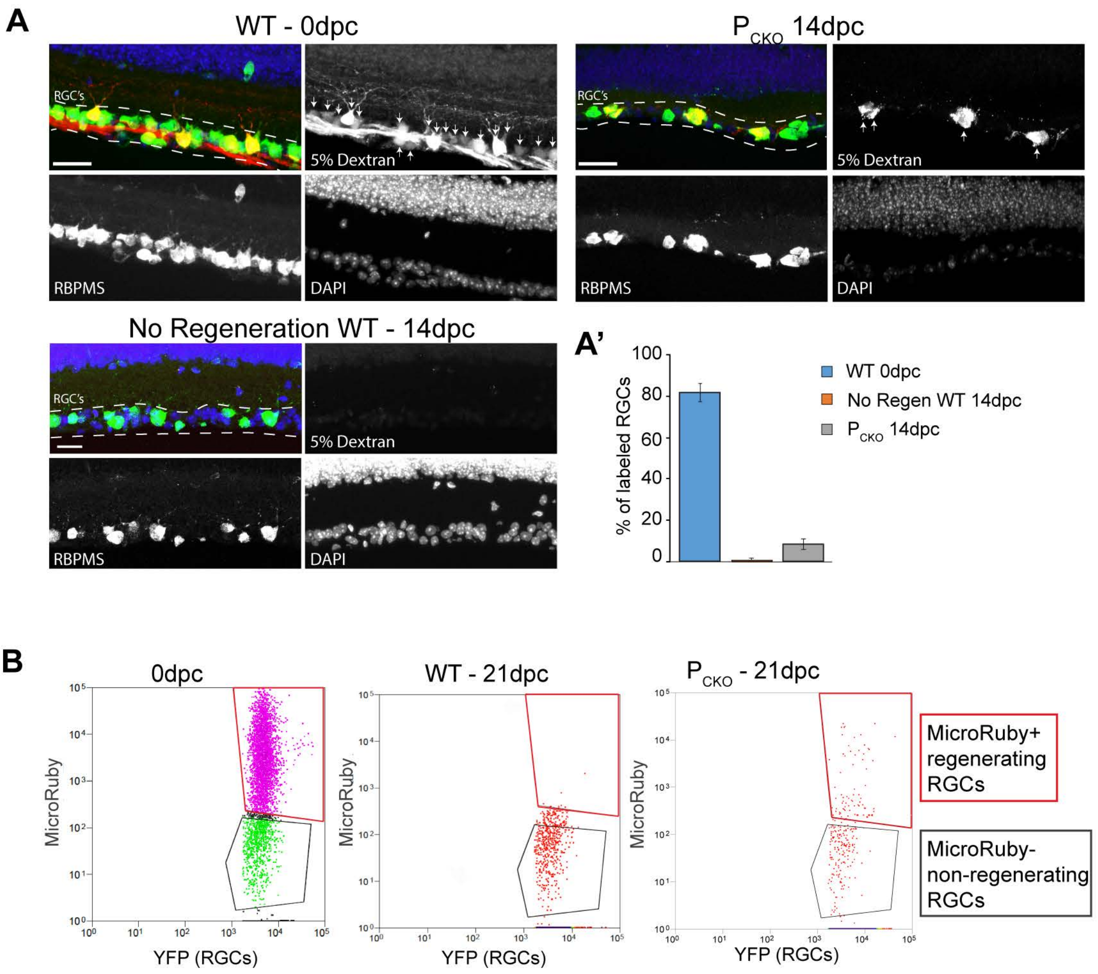

### Suppl Figure 3

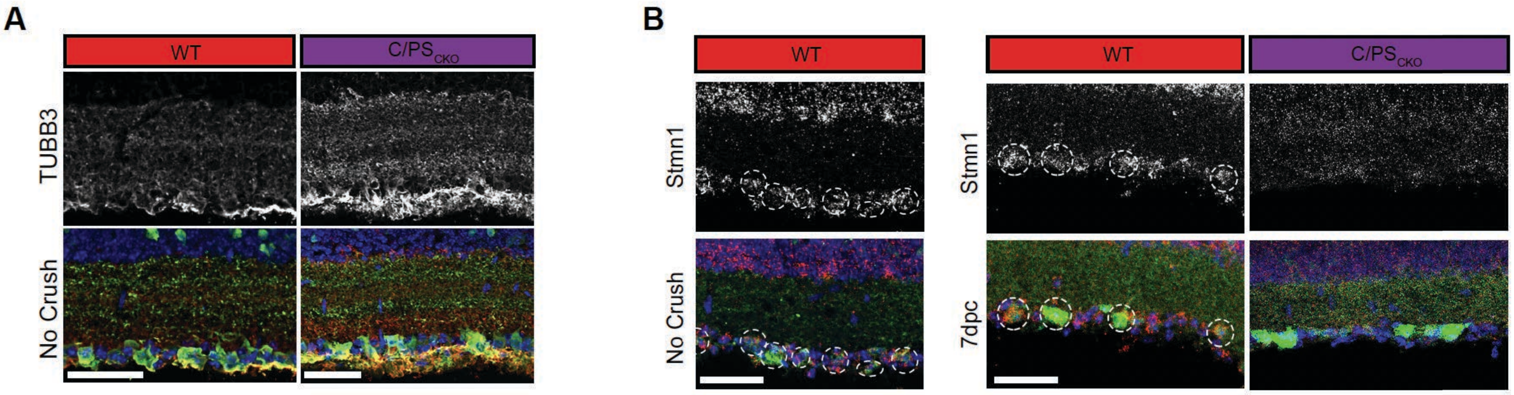


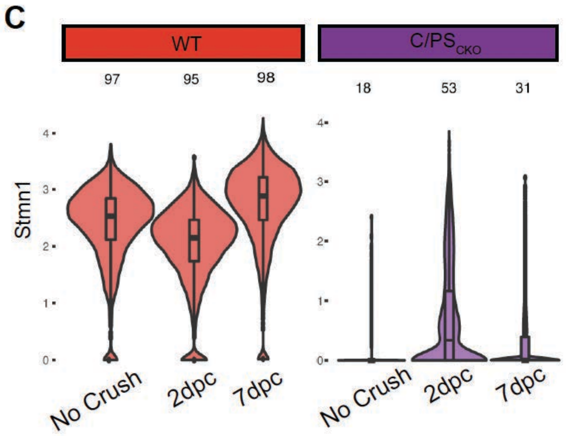

### Suppl Figure 4

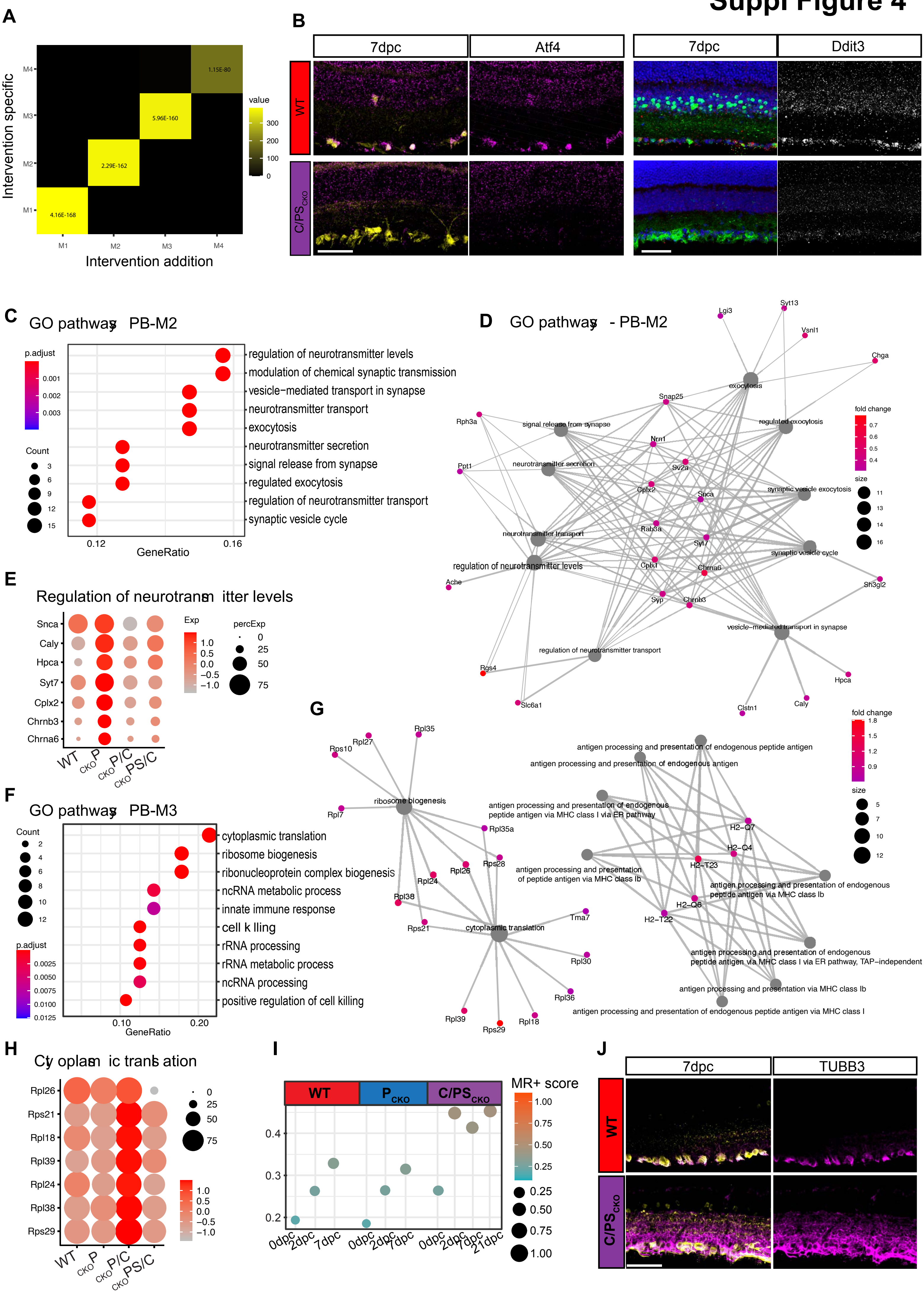

### Suppl Figure 5

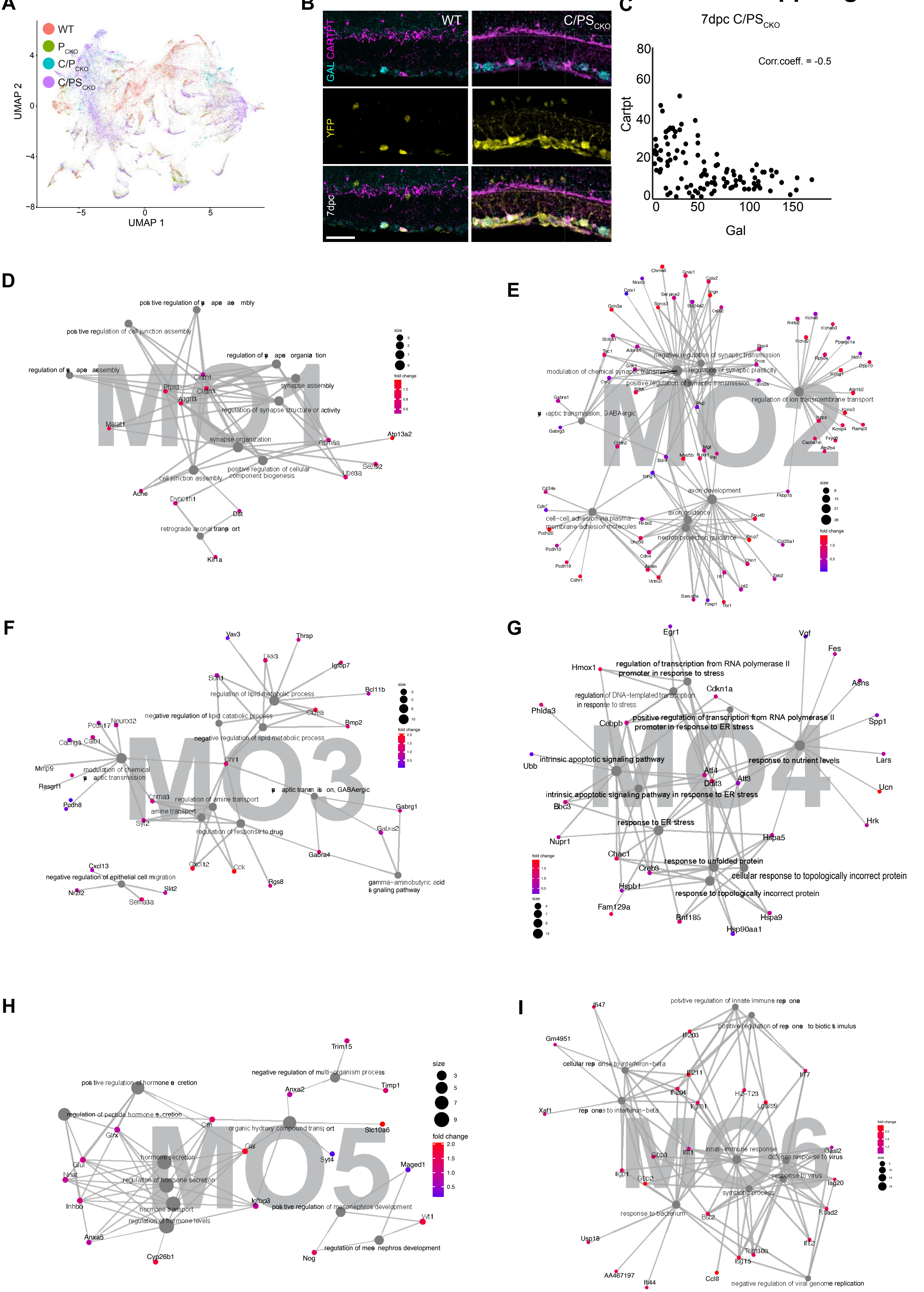

### Suppl Figure 6

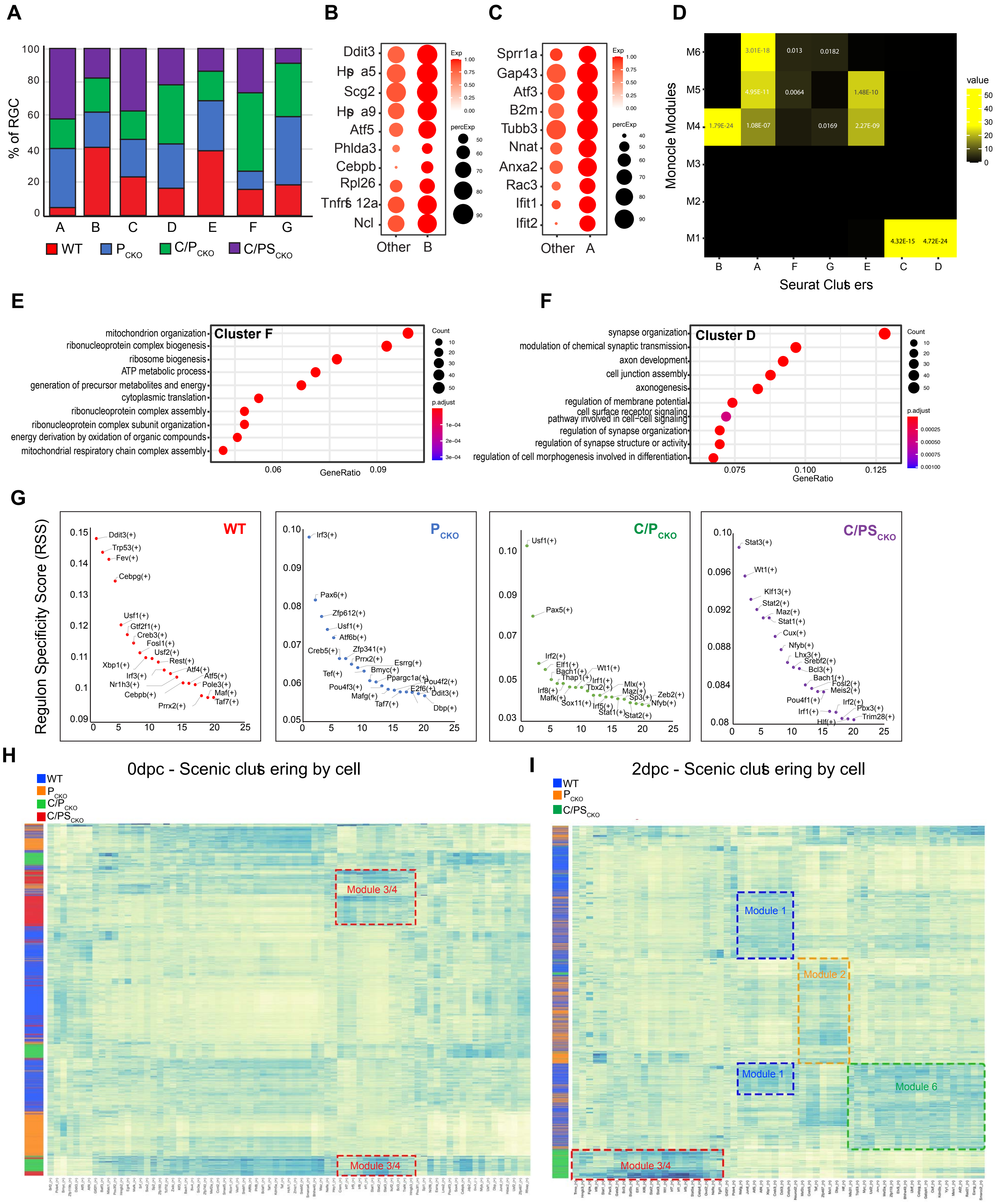

### Suppl. Figure 7

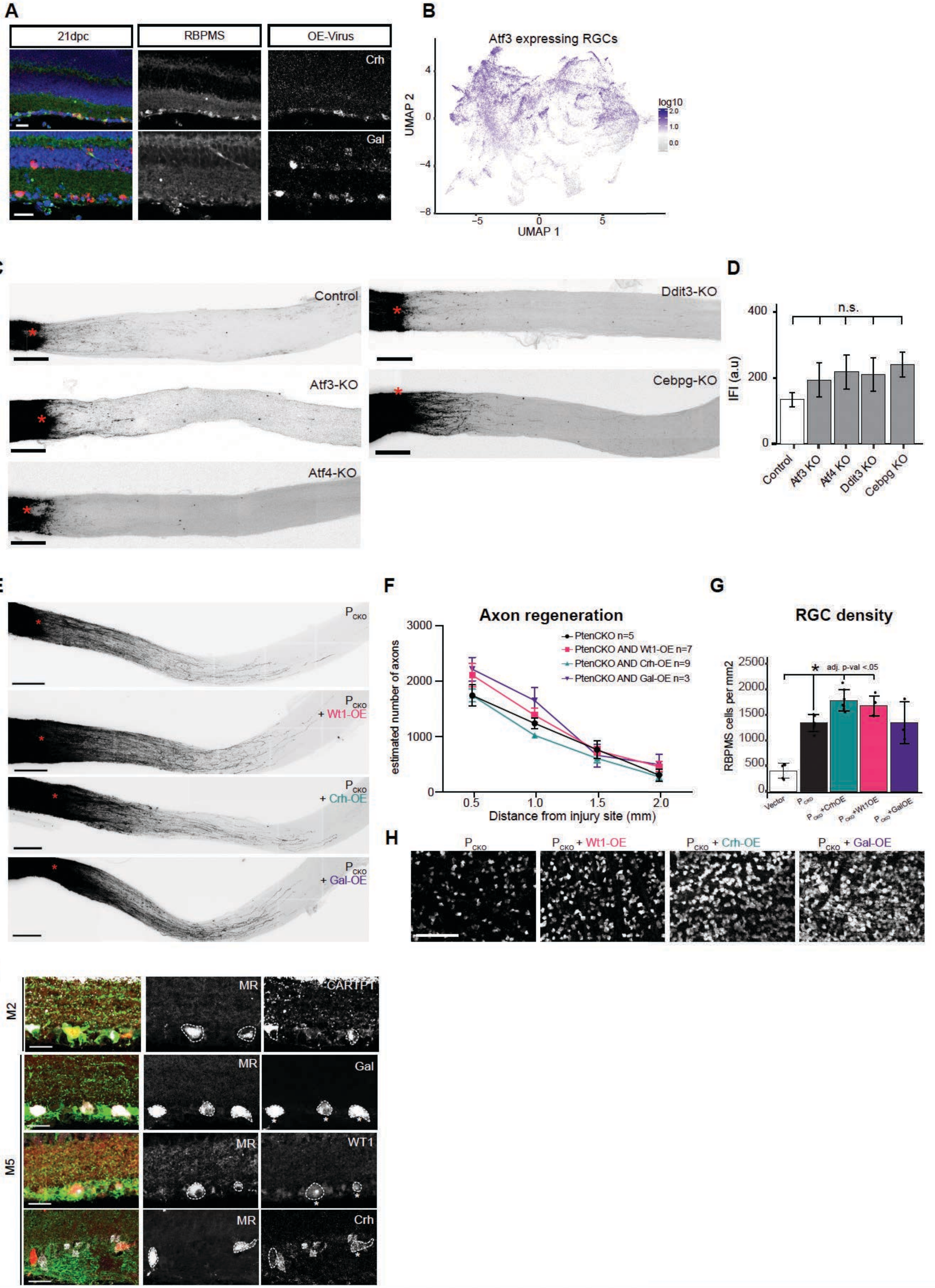
